# Human cytomegalovirus Fcγ-binding proteins constrain CD16a-driven adaptive NK cell expansion

**DOI:** 10.64898/2026.11.25.769103

**Authors:** Magdalena Huber, Niclas Schierloh, Sophie Heese, Miriam Sütterle, Patricia Otto-Mora, Günther Päth, Markus Umhau, Miguel López-Botet, Hannes Vietzen, Elisabeth Puchhammer-Stöckl, Bertram Bengsch, Hartmut Hengel, Philipp Kolb

**Affiliations:** Institute of Virology, Medical Center, Faculty of Medicine, University of Freiburg, Freiburg, Germany; Department of Medicine II, Medical Center, Faculty of Medicine, University of Freiburg, Freiburg, Germany; Signalling Research Centres BIOSS and CIBSS, Freiburg, Germany; Institute for Transfusion Medicine and Gene Therapy, University of Freiburg and Faculty of Medicine, University Hospital Freiburg, Freiburg, Germany; Hospital del Mar Medical Research Institute (IMIM). Barcelona, Spain; Department of Medicine and Life Sciences. Univ. Pompeu Fabra. Barcelona, Spain; Center for Virology, Medical University of Vienna, Vienna, Austria

## Abstract

Adaptive natural killer (NK) cells accumulate during human cytomegalovirus (HCMV) infection, but whether viral immune-evasion proteins regulate their formation is unknown. We found that the CD16a-activating capacity of HCMV-specific IgG correlated with adaptive NK cell abundance in seropositive individuals and that CD16a engagement preferentially expanded NKG2C+ NK cells. HCMV-encoded viral Fc-gamma receptors (vFcγRs) gp34 and gp68 opposed this response: virions lacking vFcγRs enhanced IgG-dependent adaptive NK cell proliferation, increased NKG2C+CD16a+ populations, and augmented degranulation after antibody-dependent stimulation, whereas gp34 or gp68 expression suppressed expansion. Flow and mass cytometry further showed that vFcγRs altered the composition and phenotypic state of expanded NK cells. Thus, HCMV vFcγRs extend immune evasion beyond acute FcγR blockade by constraining antibody-driven adaptive NK cell expansion and functional responsiveness.

## Introduction

Human cytomegalovirus (HCMV) is a beta-herpesvirus with high worldwide sero-prevalence rates of 56-94% (*1*). It establishes lifelong persistent infection, which is characterized by recurring phases of latency and reactivation. A hallmark of HCMV infection is the emergence and long-term expansion of adaptive natural killer (NK) cells, a specialized NK cell population characterized by enhanced cytolytic effector function, altered receptor repertoires, and memory-like properties (*2–7*). These cells are commonly defined by expression of NKG2C and display epigenetic and functional features distinct from conventional NK cells, including enhanced antibody-dependent cellular cytotoxicity (ADCC) (*8, 9*). Several studies have implicated HLA-E recognition by NKG2C in adaptive NK cell expansion, particularly through presentation of HCMV-derived UL40 peptides (*10–12*). However, accumulating evidence indicates that FcγR signaling is also central to adaptive NK cell biology. Adaptive NK cells exhibit enhanced responsiveness to Fc-gamma receptor (specifically FcγRIIIA/CD16a) stimulation, and genetic deficiency of CD16a prevents adaptive NK cell expansion in humans (*9, 13*). Moreover, IgG-opsonized HCMV-infected cells can selectively drive adaptive NK cell proliferation in vitro (*14*). This suggests that Fcγ-dependent receptor activation constitutes a major axis governing adaptive NK cell differentiation during HCMV infection. HCMV has evolved numerous immune-evasion mechanisms that antagonize innate and adaptive immunity to enable lifelong persistence (*15, 16*). Among these are the viral Fcγ-binding glycoproteins (viral Fc-gamma receptors, or vFcγRs) gp34 (*RL11*) and gp68 (*UL119-118*), which interfere with host FcγR activation and protect infected cells from ADCC (*15, 17, 18*). Recent work in the nonhuman primate model demonstrated that vFcγRs are critical for lytic replication of rhesus CMV (RhCMV) infection *in vivo*, underscoring their central role in CMV immune evasion from IgG-mediated humoral immunity and persistence (*19*). These proteins bind distinct regions of IgG-Fc (Fcγ) and cooperatively antagonize Fcγ-mediated immune recognition by limiting engagement of host FcγRs and removal of immune complexes from the cell surface (*15*). While this mechanism is well-established in the context of effector cell inhibition, it has remained unknown whether vFcγRs also influence the development and functional imprinting of adaptive NK cells. Here, we show that the CD16a activation potential of HCMV-specific IgG correlates with adaptive NK cell abundance in HCMV-seropositive individuals and that IgG/CD16a signaling alone is sufficient to drive preferential expansion of adaptive NK cells over conventional NK cells. We further demonstrate that the HCMV-encoded vFcγRs gp34 and gp68 suppress this process by antagonizing Fcγ-dependent receptor activation. Genetic deletion of vFcγRs enhances adaptive NK cell proliferation, promotes acquisition of a highly reactive NKG2C^+^CD16a^+^ phenotype, and augments antiviral effector function. Together, our findings reveal that HCMV immune-evasion molecules regulate not only acute Fcγ-mediated responses but also the differentiation trajectory and functional specialization of adaptive NK cells, identifying vFcγRs as modulators of NK cell differentiation.

## Results

### CD16a activation by HCMV-IgG predicts adaptive NK abundance in HCMV infection

Previous *in vitro* work revealed that immune IgG can drive adaptive NK cell expansion *in vitro* (*14*). Together with the observation that CD16a expression is required for the emergence of adaptive NK cells in CMV-infected individuals (*13*), we hypothesized that HCMV-specific IgG is sufficient to induce adaptive NK cell expansion via CD16a activation. To test this hypothesis, we incubated HCMV (TB40/E) infected MRC-5 fibroblasts with titrated sera from 20 HCMV-seropositive and 5 HCMV-seronegative healthy individuals and probed them for CD16a activation using a cell-based reporter assay. In brief, we co-cultured the opsonized infected cells with BW5147 cells expressing human FcγRs and quantified reporter cell activation by mIL-2 ELISA (*20*) (Fig. 1A). Adaptive NK cells were identified as NKG2C^+^/CD3^-^/CD56^+^ (Fig. S1A), with CD16a as an additional marker, which had no significant influence on adaptive NK cell frequency determination in individuals (Fig. S1B). In the human population, individuals can differ regarding the expression of NKG2C or the two major CD16a isoforms. It has been reported that individuals with heterozygous V/F or homozygous V/V expression of the 158V variant have a higher reactivity compared to the F/F variant (*21*). Our cohort was genotyped for individual NKG2C and CD16a genotypes (Fig. 1B). Two individuals had a homozygous deletion for NKG2C and were therefore excluded due to our adaptive NK cell definition from further analysis. We found that V/F and V/V genotypes generally coincide with a higher adaptive NK cell frequency, supporting the conclusion that differences in CD16a reactivity impact adaptive NK cell expansion (Fig. 1C). As not all HCMV-seropositive individuals develop a substantial population of adaptive NK cells (*22*) (Fig. S1B), we used data generated from the HCMV-seronegative group to define a threshold for increased adaptive NK cell frequency. This revealed a significant correlation between an individual’s adaptive NK cell frequency and the CD16a activation by their respective anti-HCMV IgG (Fig. 1D) [p=0.02139, r=0.7667]. Fittingly, when we exclude CD16a reactivity from the analysis, we find no significant correlation between the overall anti-HCMV IgG titer alone and adaptive NK cell frequency (Fig. 1E) [p=0.2696, r=0.4176]. For comparison, CD16a activation was also mediated by EBV IgG upon opsonisation of EBV-infected P3HR1 cells, representing another latently infecting herpesvirus with high seroprevalence, which did not result in a significant correlation with adaptive NK cell frequency (Fig. 1F) [p=0.8801, r=0.006667], anti-EBV IgG titers alone (Fig. 1G) [p=0.2696, r=-0.45] or human FcγRI/CD64 activation by anti-HCMV IgG (Fig. 1H) [p=0.2696, r=0.4167]. This again confirms the specific involvement of CD16a in adaptive NK cell expansion in HCMV seropositive individuals, but also implies further unique features of HCMV-specific IgG to impact CD16a mediated adaptive NK cell expansion in a subset of HCMV seropositive individuals.

**Fig. 1.**
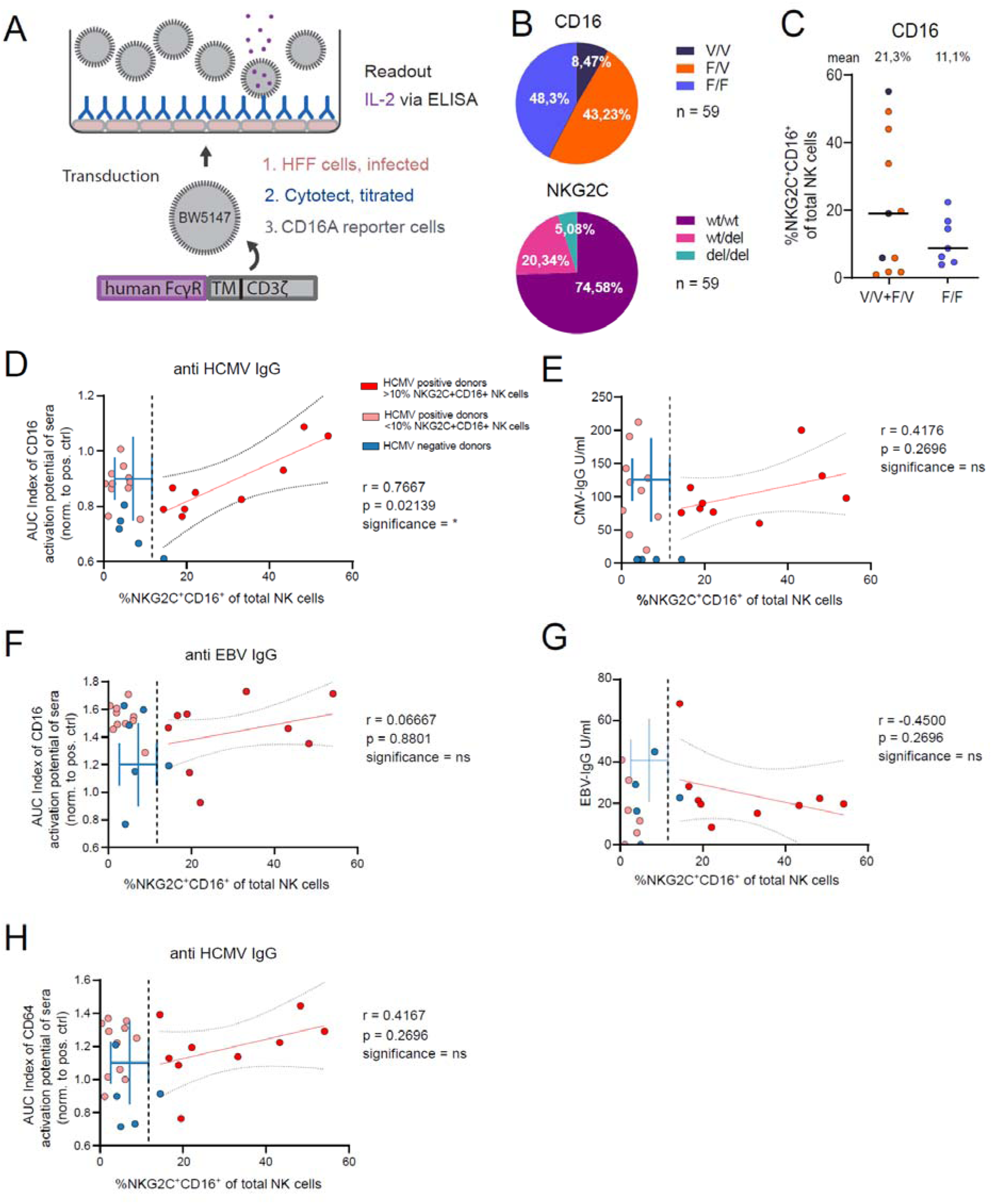
HCMV-specific CD16a activation correlates with adaptive NK cell abundance in humans. **(A)** To determine FcγR activation potential by HCMV-specific or EBV-specific IgG from seropositive individuals a cell-based reporter assay was used. MRC-5 cells were infected with HCMV TB40/E and incubated with titrated amounts of serum. Opsonized cells were then co-cultured with BW5147 reporter cells stably expressing human FcγR ectodomains genetically fused to the mouse CD3ζ signaling module. Receptor activation results in mIL-2 secretion, which was quantified via ELISA and is expressed as an area under curve (AUC) index value. **(B)** CD16a and NKG2C genotypes across cohort. **(C)** Rates of NKG2C^+^/CD16a^+^ NK cells grouped by CD16a genotype. **(D-H)** NKG2C^+^/CD16a^+^ NK cell rates of CMV positive (n=20) and CMV negative (n=5) donors were correlated with **(D)** CD16a activation potential by anti-HCMV IgG, **(E)** serum anti-HCMV IgG titers, **(F)** CD16a activation potential by anti-EBV IgG, **(G)** anti-EBV IgG titers or **(H)** CD64 activation potential by anti-HCMV IgG. A threshold of NKG2C^+^ NK cell occurrence was set according to NKG2C^+^ NK cell frequencies measured for HCMV seronegative donors (Mean = 7.132%, SD = 4.5, total threshold = 11,63%). Spearman correlation, plotted 95% confidence interval.

### CD16a activation selectively drives adaptive NK cell expansion over conventional NK cells

Given that adaptive NK cells express a more reactive CD16a/CD3-ζ complex compared to conventional CD16a/FcεRγ-chain expressing NK cells^12,18^ and following the observation that adaptive NK cell frequency correlates with CD16a activation by anti-HCMV IgG *in vivo*, we tested the impact of CD16a activation on adaptive NK cell expansion specifically in an *in vitro* setting of reduced complexity. To this end, we designed an assay platform that allows for the surveillance of cellular proliferation and expansion over time (Fig. 2A). In brief, PBMCs from HCMV-seropositive (n=19) or –seronegative (n=11) individuals were cultured on immobilized human/humanized IgG (trastuzumab/Herceptin® or rituximab) for 7 days to elicit CD16a activation. Subsequently, the proliferation of stimulated NK cells was evaluated. Endpoint analysis via flow cytometry included staining for CD3, NKG2C, CD56 and CD57 to identify conventional and adaptive NK cells as well as T cells, which do not typically express FcγRs. The gating strategy is shown in Fig. S1A. Other than IgG, no additional stimuli were added. An IgG dose-dependent increase in proliferation was detected for CD3^-^/CD56^+^ NK cells but not for CD3^+^ T cells (Fig. 2B and Fig. S1C). Compared to the total lymphocyte count, NKG2C^+^ adaptive NK cells showed increased proliferation over NKG2C^-^ conventional NK cells. When comparing seropositive/-negative individuals regarding matched IgG-mediated NKG2C^+^ NK cell expansion, we found that NKG2C^+^ NK cell expansion is significantly enhanced compared to NKG2C^-^ cells, but only regarding NK cells from seropositive individuals (Fig. 2C). This increased frequency of NKG2C^+^ NK cell expansion in seropositive individuals was strictly IgG dependent and not seen for NKG2C^-^ NK cells or T cells (Fig. 2D). Conversely, utilizing the same assay setup we then tested if adaptive NK cell expansion is also strictly CD16a dependent. To this end, we assessed NKG2C^+^ NK cell expansion in a representative donor on immobilized human IgG and in the presence of titrated amounts of FcγR-specific blocking monoclonal antibodies (mAbs, clone 3G8). We found that equimolarity between immobilized IgG and 3G8 completely abrogates expansion and antagonizes proliferation, which suggests strict CD16a dependency (Fig. S2A). Conversely, expansion and proliferation by titrated amounts of immobilized 3G8 (Fig. S2B) or immobilized recombinant human IgG (Fig. S2C) was nearly identical, further substantiating this conclusion. A reduced cell count resulting from the use of high amounts of immobilized IgG implies overstimulation, as proliferation was impaired in the later phases of expansion. IgG-dependent NKG2C^-^ NK cell expansion also showed strict CD16a dependency but at a generally lower level (Fig. S2A-C), in line with the above observation that NKG2C^+^ NK cells are more reactive towards immobilized IgG compared to NKG2C^-^ NK cells (Fig. 2B). We then extended these experiments to include data from a total of 11 HCMV-seropositive individuals at equimolarity as in Fig. S2A-C. Primary data for an exemplary donor is shown in Fig. S2D. Remarkably, stimulation by immobilized IgG was completely abolished when 3G8 was added to block CD16a, clearly showing that CD16a activation is sufficient to drive expansion of NKG2C^+^ NK cells. Of note, immobilized 3G8 was able to stimulate expansion of NKG2C^+^ NK cells, albeit at lower levels compared to immobilized IgG (Fig. S2E).

**Fig. 2.**
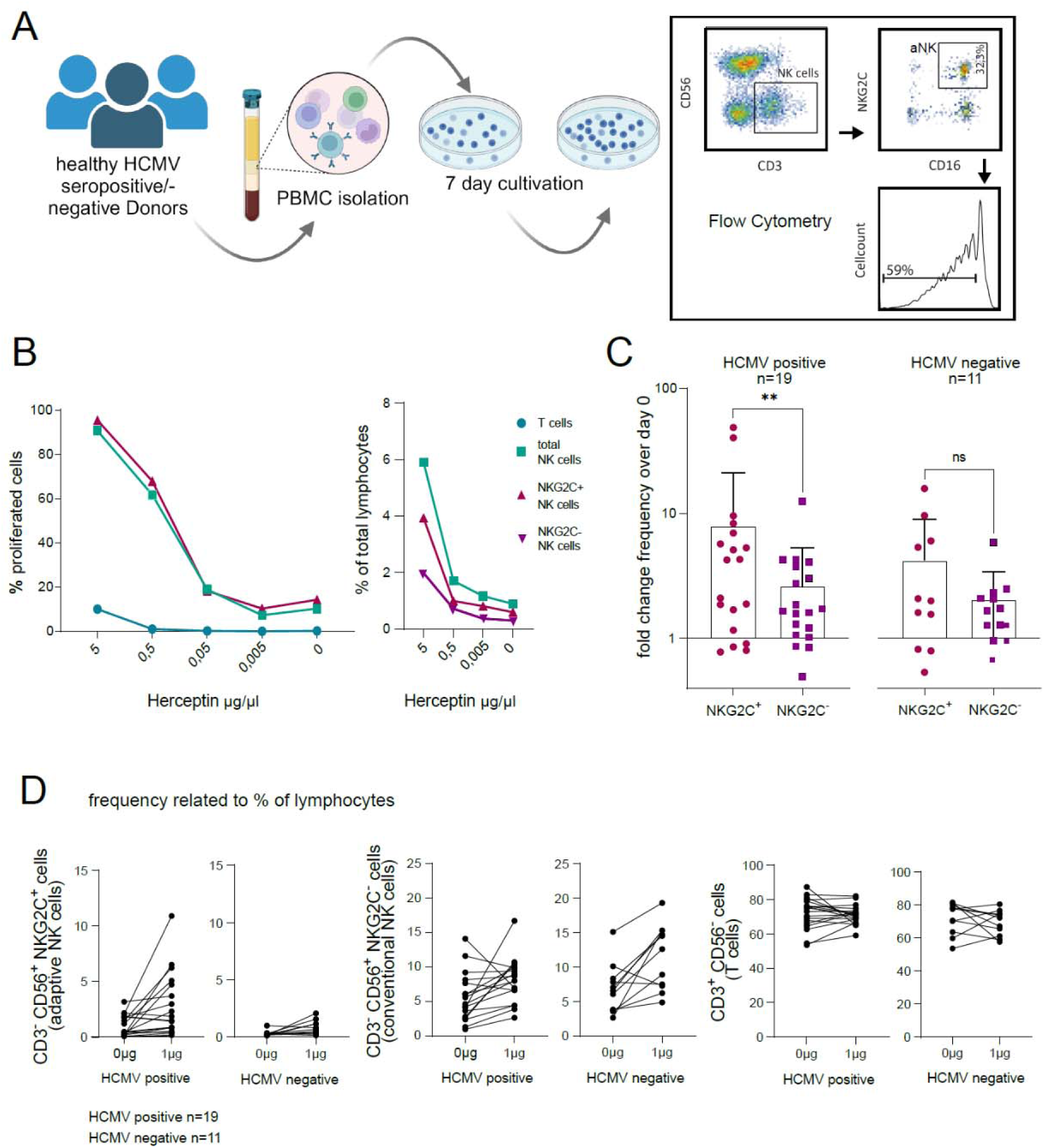
CD16a activation selectively drives adaptive NK expansion. **(A)** Scheme of the IgG based expansion assay: PBMCs of healthy donors were pre-stained with proliferation dye and cultured on immobilized human IgG. After 7 days of incubation, percentage and proliferation rate of immune populations were acquired via flow cytometry. **(B)** Titrated amounts of anti-HER2 IgG1 (trastuzumab/Herceptin®) were used to induce FcγR activation. Percentage of proliferated cells and percentage of NK cells (CD3^-^ CD56^+^), adaptive NK cells (CD3^-^ CD56^+^ NKG2C^+^), and T cells (CD3^+^) in relation to total lymphocytes are shown. Mean values of 2 experiments. One exemplary donor. **(C)** Fold change of NKG2C^+^ or NKG2C^-^ NK cell frequencies at day 7 over day 0 comparing HCMV sero-positive and –negative individuals (n=19). **(D)** Frequencies of adaptive NK cells in total lymphocytes from HCMV sero-positive (n=19) and -negative donors (n=11) in the presence [1µg] and absence [0µg] of immobilized human IgG1 (rituximab). ***=p<0.0002,**=p<0.0021, *=p<0.0332, (Wilcoxon test, two tailed).

### HCMV encoded vFcγRs counteract FcγR-dependent adaptive NK cell expansion

As we found IgG-mediated adaptive NK cell expansion to be CD16a dependent, we tested whether HCMV vFcγRs affect CD16a-mediated adaptive NK cell expansion. The HCMV virion envelope displays all major HCMV antigens found on the surface of an infected cell, including HCMV vFcγRs (*23–26*). We therefore introduced HCMV virions into the above assay platform to assess adaptive NK cell expansion in the context of a natural HCMV specific IgG response. Specifically, we immobilized titrated amounts of virions derived from the HCMV AD169 strain, lacking the pentameric entry complex essential for the infection of certain cell types (*27*), or a vFcγR-deleted (Δgp34 [*RL11*], Δgp68 [*UL119-118*] and Δgp95 [*RL12*]) variant thereof, termed AD169ΔΔΔ (*15*) and incubated them with fixed concentrations of hyper immunoglobulin (HIG) containing HCMV specific IgG (Cytotect^®^) (Fig. 3A). We also tested the endotheliotropic TB40/E strain for comparison (*28*). Comparing binding of HIG to immobilized virions of different HCMV strains/mutants showed comparable amounts of opsonizing IgG (Fig. S3A). NKG2C^+^ adaptive NK cell expansion depended on the number of immobilized virions and was comparable between HCMV AD169 and TB40/E virions but enhanced on vFcγR deleted AD169 virions (Fig. 3B), which was not observed for NKG2C^-^ NK cells (Fig. 3C). Accordingly, NKG2C^+^ NK cells showed stronger and earlier formation of lymphoblasts compared to NKG2C^-^ NK cells (Fig. 3D). NKG2C^+^ NK cell expansion on a fixed number of immobilized virions and incubated with titrated amounts of HIG lead to comparable results with stronger expansion (Fig. 3E) and proliferation (Fig. 3F) on vFcγR-deleted virions and at lower concentrations of IgG. Comparing expansion on titrated amounts of immobilized virions in the presence or absence of HIG clearly shows that HCMV vFcγRs limit expansion in an IgG-dependent manner (Fig. 3G). Comparing the NKG2C^+^ NK cell frequencies following 7 days of culture on virions of full-length AD169, mouse cytomegalovirus (MCMV) or PBS demonstrated a clear requirement for HCMV virions to provide binding to target antigens for opsonization by IgG (Fig. S3B). We also confirmed that the observed vFcγR-mediated effects on NKG2C^+^ NK cell expansion and proliferation were not limited to the use of opsonized virions but was recapitulated using opsonized infected cells (Fig. S4A). Testing NKG2C^+^ NK cell expansion with opsonized infected cells of AD169 mutants expressing gp34 or gp68 individually and in the absence of gp95 (*15*), revealed an effect of gp68 over gp34 in this setting (Fig. S4B). As gp68 is known to interfere with CD16a binding to immune complexes directly (*15*), this suggests that the block of immune complex engagement by CD16a is the leading mechanism by which HCMV controls IgG-mediated NKG2C^+^ NK cell expansion. To test whether HCMV vFcγRs are able to limit IgG-dependent adaptive NK cell expansion in the absence of infection, we recombinantly expressed gp34 or gp68 or both in 293T cell stably expressing CD20, which served as a target antigen for the humanized IgG1 antibody rituximab (RTX) (Fig. S4C). Here, in the absence of other viral factors, we again find that IgG-dependent NKG2C^+^ NK cell expansion is antagonized by both vFcγRs individually or in combination. We next tested if cellular expression of vFcγRs is required to limit NKG2C^+^ NK cell expansion by adding recombinantly produced and purified ectodomains of HCMV vFcγRs to SKOV3 cells stably expressing the HER2 antigen and opsonizing them using trastuzumab (human IgG1) (Fig. S4D). This showed that the ectodomains of gp34, gp68 or a combination of both are sufficient to counteract expansion. As the ectodomains of HCMV vFcγRs do not mediate immune complex internalization in this setting, this again confirmed that the block of immune complex engagement by CD16a is sufficient to achieve the effect. Of note, probing with a non-Fcγ-binding point mutant of gp34, gp34mtrp (*15*), shows that Fcγ binding is essential for HCMV vFcγRs to limits IgG-dependent NKG2C^+^ adaptive NK cell expansion.

**Fig. 3.**
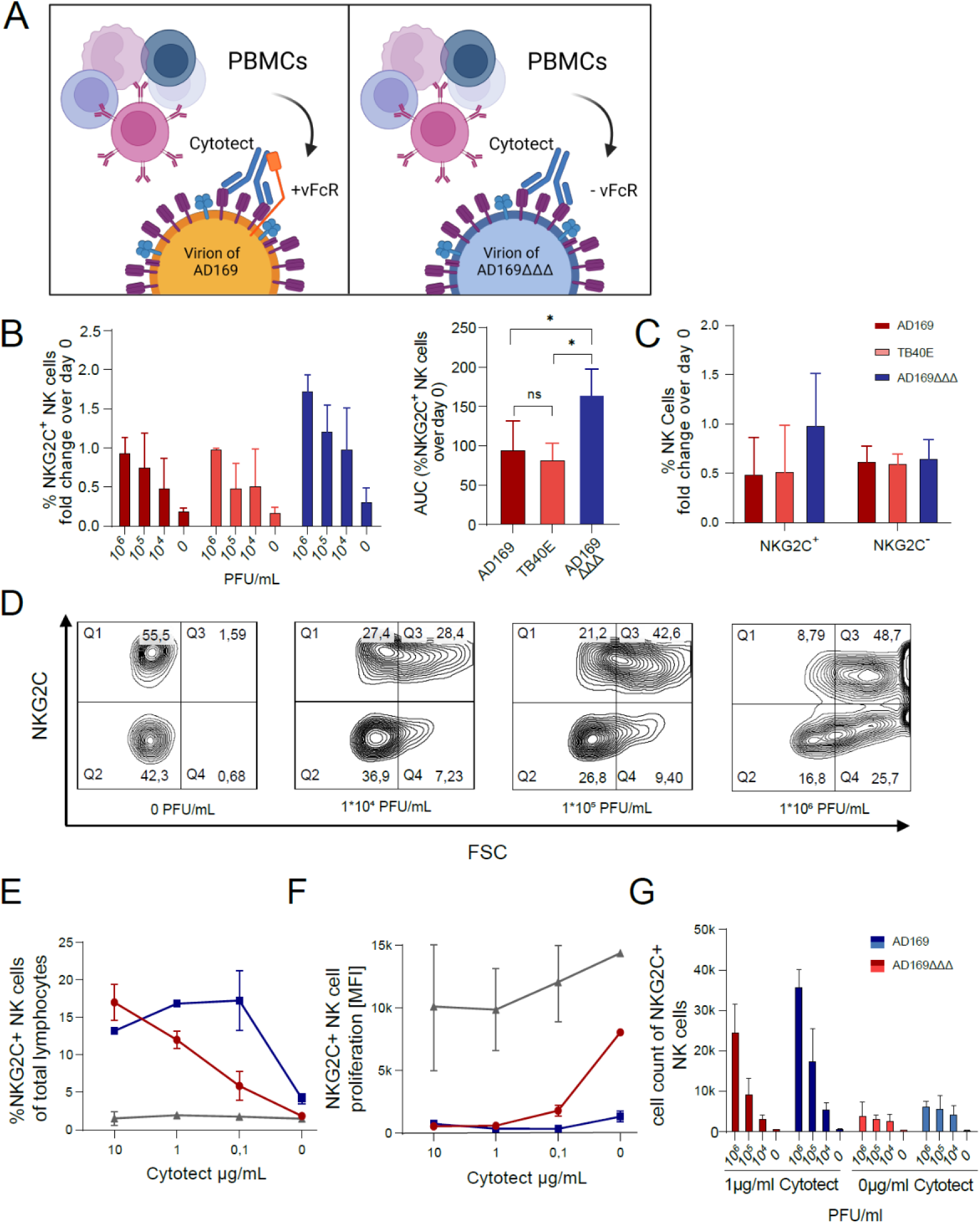
HCMV vFcγRs limit FcγR-dependent adaptive NK cell expansion. **(A)** Scheme of the virion-based expansion assay. HCMV virions were immobilized and incubated with HIG containing HCMV-specific IgG (Cytotect®). PBMCs were then cultured on immobilized and opsonized virions for 7 days followed by flow cytometry analysis. Equal amounts of PBMCs were analysed in a 60 s time gate and normalized to day 0. **(B)** NKG2C^+^ NK cell expansion comparing culture on titrated amounts of immobilized HCMV virions containing (AD169 and TB40/E) or lacking vFcγRs (AD169ΔΔΔ). Virions were opsonized using 50µg/ml HIG. Right graph shows area under curve (AUC) evaluation of the titration. Two independent experiments performed in technical replicates. (SD = range, *=p<0.05, 1-way analysis of variance (ANOVA), Fishers LSD). **(C)** Comparison between NKG2C^+^ and NKG2C^-^ NK cells at 1*10^4^ PFU. Two independent experiments performed in technical replicates. SD = range. **(D)** Density plots of the total NK cell population differentiated by NKG2C expression and following culture on titrated amounts of opsonized AD169 virions show asynchronous NK cell blasting. Gating on CD3^-^/CD56^+^ cells. Q1 shows non-blasting adaptive NK cells (NKG2C^+^/FSC^-^), Q2 shows blasting adaptive NK cells (NKG2C^+^/FSC^+^), Q3 shows non-blasting NK cells (NKG2C^-/^FSC^-^) and Q4 shows blasting NK cells (NKG2C^-^ /FSC^+^). Bar graph shows percentage of 3 independent experiments. **(E, F)** 1*10^5^ PFU were immobilized as above and opsonized using HIG as indicated. Percentage of NKG2C^+^ NK cells following 7-day expansion in relation to total lymphocytes were analysed for cell count **(E)** and proliferation **(F)**. **(G)** Graded amounts of virions were immobilized and opsonized using 0µg or 10µg of HIG before NKG2C^+^ cell counts were measured.

### HCMV vFcγRs limit acquisition of highly reactive adaptive NK subsets

We speculated that limiting the expansion of highly antiviral adaptive NK cells by vFcγRs is an intended surplus effect of vFcγR-mediated protection from ADCC. Further, we hypothesized that restriction of adaptive NK cell expansion impacts the composition of the NK cell compartment. To test this assumption, we cultured PBMCs from 10 HCMV seropositive individuals on immobilized AD169 virions opsonized by HIG (Cytotect^®^) as above. NK cells were characterized via flow cytometry comparing subpopulations before and after IgG-dependent NK cell expansion. This confirmed that IgG-dependent adaptive NK cell expansion is limited in the presence of HCMV vFcγRs across HCMV seropositive donors (Fig. 4A). High-dimensional analysis of a representative donor illustrated that the absence of vFcγR expression leads to an expansion of predominantly NKG2C^hi^/CD16a^hi^ NK cells (Fig. 4B). Of note, FcεRγ downregulation was previously shown to be associated with increased IgG-mediated proliferation *in vitro* (14), however, we found NKG2C-positivity to be a more precise marker of expanding NK cells (fold change) (Fig. S5A). Overall, CD16a dependent expansion gives rise to a dominant NKG2C^hi^/CD16a^hi^ population of adaptive NK cells (Fig. S5B), in line with our above observations of strict CD16a/IgG-dependence and the observation that CD16a reactivity of CMV-specific IgG correlates with adaptive NK cell frequency in humans (Fig. 1D). We then performed population analyses on merged data from all 8 donors following IgG-dependent expansion using FlowSOM (*29*). This analysis revealed that compared to day 0, a prominent NKG2C^hi^/CD16a^hi^/NKG2A^lo^ population forms upon expansion on opsonized virions lacking vFcγR expression (Fig. 4C). Additionally, we find that FcεRγ negativity only coincides with IgG-dependent expansion in certain adaptive NK cell populations, while NKG2C-positivity consistently coincides with increased CD16a expression on more NK subgroups (Fig. S5B-C). Conversely, NKG2C^+^CD16a^+^ NK cells appear at a lower rate (WT+IgG: 25,4% versus ΔΔΔ +IgG: 38,7%) on vFcγR-containing virions and generally remains at a low rate in the absence of IgG regardless of vFcγR expression (Fig. 4D).

**Fig. 4.**
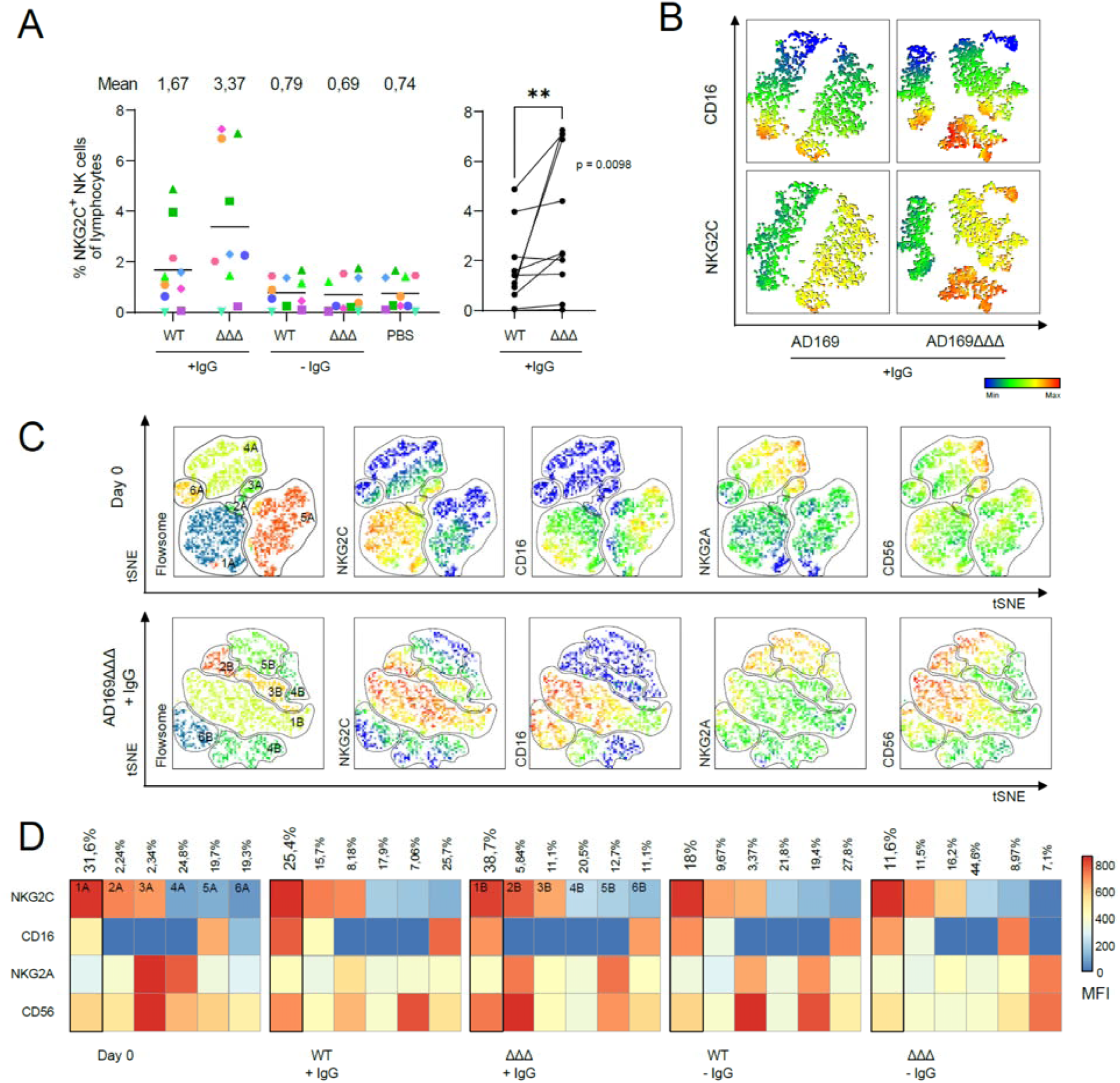
vFcγRs limit formation of highly reactive adaptive NK cell populations. PBMCs from HCMV positive healthy donors (n=10) were cultured for 7 days on immobilized virions (1*10^6^ PFU/ml) containing (AD169) or lacking (AD169ΔΔΔ) vFcγRs. Virions were opsonized using 10µg HIG or not. Equal amounts of PBMCs were analysed in a 60 s time gate. **(A)** Percentage of NKG2C^+^ NK cells compared to total lymphocyte counts (left) and paired comparison of NKG2C^+^ NK cells between opsonized AD169 and AD169ΔΔΔ (right). *=p<0.05, two tailed Wilcoxon matched-pairs signed rank test. **(B)** NK cell analysis of one representative using tSNE high dimensional analysis to compare CD16a and NKG2C expression upon NK cell expansion on opsonized AD169 or AD169ΔΔΔ virions. **(C)** Pooled analyses as in **(B)** integrating data from 8 donors before and after expansion on opsonized virions. Before data integration, counts were set to 1.000 cells to ensure equal representation. Population demarcation was performed using FlowSOM and populations were analysed for expression of NKG2C, CD16a, NKG2A and CD56. Heat map shows rates of all populations classified via FlowSOM before (d0) and after cultivation as in **(A)**. Color scale indicates MFI.

### HCMV vFcγRs imprint adaptive NK cell function

As we found that vFcγRs affect the IgG-dependent expansion of adaptive NK cells resulting in a changed composition of the NK cell subpopulations, we assessed how this shift within the pool of NK cells affects the overall effector responsiveness. To test the effector function after IgG mediated expansion in presence or absence of vFcγRs, we compared PBMCs from select donors (n=5) for their degranulation potential after culture on opsonized wildtype or vFcγR-deleted AD169 virions. Following co-culture with opsonized virions as above (AD169 versus vFcγR-deleted variant AD169ΔΔΔ), PBMCs were rested for 12 – 24 hr. Subsequently, degranulation was assessed by exposure to 1 µg or 0 µg immobilized IgG as a stimulus to exclude additional receptor-ligand interactions. Adaptive NK cells defined as NKG2C^+^CD16a^+^ were compared to subpopulations that were double-negative for both markers (NKG2C^−^CD16a^−^), and to cells that were single positive for each marker (NKG2C^+^CD16a^−^ and NKG2C^−^CD16a^+^). Among PBMCs expanded with opsonized AD169ΔΔΔ, the frequency of CD107a^+^ NK cells responding to IgG was highest in the adaptive NK cell population, surpassing other subpopulations (Fig. 5B). In contrast, when cells were expanded with AD169, where vFcγR expression diminishes IgG-mediated expansion, a reduction in CD107a^+^ NK cells was observed. The same trend was noted for the mean fluorescence intensity (MFI) of CD107a, indicating a higher expression level, and thus greater reactivity, within subpopulation (Fig. 5C). While NKG2C^−^CD16a^+^ single-positive NK cells also degranulated following IgG stimulation, their degranulation levels were lower than those of adaptive NK cells expanded in the absence of vFcγRs. Additionally, a 7-day long-term culture without opsonized virions or other stimuli resulted in lower degranulation levels in adaptive NK cells compared to those of single expression NKG2C^-^CD16a^+^. This suggests that IgG-mediated expansion in the absence of vFcγRs drives a shift that enhances the reactivity of adaptive NK cells and thereby altering reactivity in the NK cell compartment, presumably due to transcriptomic changes. Moreover, a correlation analysis revealed that the magnitude of degranulation elicited by IgG within the whole NK cell compartment directly correlated with the CD16a^+^/NKG2C^+^ adaptive NK cell frequency (Fig. 5D). This finding indicates that the IgG specific antibody-dependent degranulation within the total NK cell population is reflected by an increased presence of adaptive NK cells. When performing high-dimensional analysis on cells before and after degranulation, most reactive NK cell populations were found to express CD107a alongside with NKG2C and CD16a, supporting the conclusion that adaptive NK cells are the main reactive population following expansion (Fig. 5E). This again implies that enhanced degranulation potential following expansion is not only dependent on adaptive NK cell frequency but also adaptive responsiveness of individual cells. Furthermore, the responsiveness of adaptive NK cells co-cultured with infected cells was tested to better replicate the effector-target interaction as it occurs *in vivo*. For this purpose, MRC-5 cells were infected with AD169ΔΔΔ and incubated either with or without IgG to assess IgG-dependent responsiveness in an unbiased manner. Similar to the previous experimental setup, adaptive NK cells expanded in the presence of AD169ΔΔΔ virions displayed increased responsiveness compared to those expanded on AD169. To determine IgG-specific degranulation, the frequency of CD107a^+^ adaptive NK cells generated by target cells incubated with 10µg was normalized to the negative control without IgG, thereby ruling out stimulation by other ligands expressed on infected cells. This resulted in an increased IgG-specific responsiveness in adaptive NK cells expanded in the absence of vFcγRs compared to those that were not stimulated during long-term incubation. Strikingly, PBMCs that were cultured on wildtype AD169 virions showed a lower reactivity than untreated PBMCs. This confirmed that vFcγRs not only limit adaptive NK cell expansion but generally lower reactivity of adaptive NK cells, likely on a transcriptomic level (Fig. 5F).

**Fig. 5.**
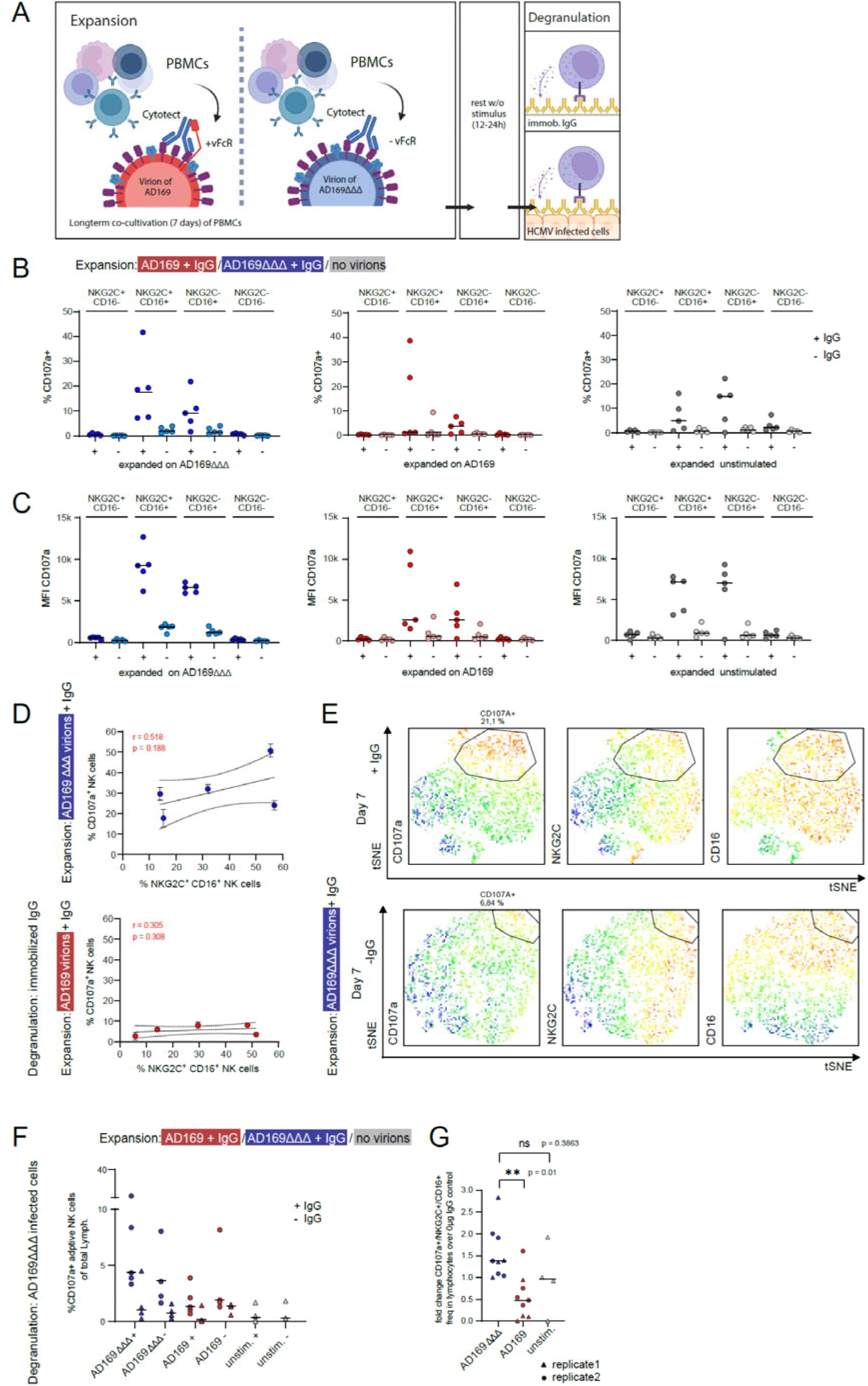
vFcγRs suppress adaptive NK cell effector responsiveness. **(A)** Scheme of the HCMV virion based *in vitro* expansion with a subsequent degranulation assay. HIG containing HCMV-specific IgG (Cytotect®) was used to opsonize HCMV AD169 or AD169ΔΔΔ virions (1*10^6^ PFU/ml) on a 0,32 cm^2^ dish and followed by co-culturing with PBMCs for 7 days (n=5). PBMCs were harvested and cultured without stimulus for 12 - 24h to rest. NK cell degranulation was induced by incubation on 1µg or 0µg immobilized IgG (shown in **B-E**) or by incubation on AD169 infected MRC-5 cells (MOI=3) **(F)** or AD169ΔΔΔ infected MRC-5 cells **(G)** opsonized using 10µg or 0µg of HIG for 5h. CD107a was used as marker to assess the level of degranulation. MFI of CD107a **(B)** and % of NK cells expressing CD107a **(C)** in NKG2C^+^/- CD16a^+^/- subpopulations following expansion in the indicated settings on the top. **(D)** Correlation of % CD16a^+^NKG2C^+^ NK cells after expansion on either opsonized AD169 (top panel) or AD169ΔΔΔ (bottom panel) virions; % cells were compared to % of total CD107a^+^ NK cells after stimulation on 1µg immobilized IgG (pearson correlation, CI 95%). **(E)** t-SNE high-dimensional analysis of NK cells from a representative donor following cultivation on opsonized AD169ΔΔΔ virions for 7 days and after degranulation on immobilized IgG or not. % of CD107^+^/NKG2C^+^/CD16a^+^ NK cells and fold change of [% of CD107^+^/NKG2C^+^/CD16a^+^ NK cells] with 10µg HIG over 0µg HIG on opsonized AD169-infected cells **(F)** or opsonized AD169ΔΔΔ-infected cells **(G)** following expansion under the conditions indicated above. Symbols show mean values from independent experiments (unpaired One Way Anova/ Kruskal-Wallis test, *=p<0.05).

### HCMV vFcγRs imprint adaptive NK cell phenotype

To determine whether the vFcγR-dependent functional differences were accompanied by phenotypic remodeling, we analyzed PBMCs from HCMV-seropositive donors by mass cytometry after 7 days of culture with IgG-opsonized AD169 virions that either expressed (+vFcγRs) or lacked (-vFcγRs) gp34 and gp68. Gaussian-parameter cleanup and combinatorial palladium barcoding were used to remove acquisition artifacts and deconvolute samples, respectively (Fig. S6). UMAP analysis resolved 12 NK cell clusters with distinct distributions of CD16, CD56, IL-2R, NKG2D, CD25, and EdU incorporation (Fig. 6A, Fig. S7). Both day-7 cultures differed from the day-0 population, but the absence of vFcγRs produced the clearest compositional shift: clusters 4, 9, 11, and 12 occupied a larger fraction, whereas clusters 2, 3, and 8 occupied a smaller fraction than in cultures containing vFcγRs (Fig. 6B). Together with cluster 10, clusters 4, 9, 11, and 12 showed a broadly similar activated and differentiated phenotype and were preferentially represented in the -vFcγR condition, consistent with enhanced responsiveness to IgG-opsonized virions in the absence of vFcγRs. Within this group, clusters 9, 11, and 12 displayed a proliferative phenotype with prominent EdU incorporation, consistent with cells in S phase, whereas clusters 4 and 10 shared much of this activated phenotype without prominent EdU incorporation. Across these clusters, the phenotype was characterized by expression of activation and differentiation-associated markers and transcription factors, including CD2, IL-2R, CD25, PLZF, T-bet, EOMES, BLIMP-1, Runx3, and mTOR. In clusters 4, 10, and 11, the -vFcγR condition was also associated with relatively higher expression of CD158a, PLZF, T-bet, EOMES, FcεR1, BLIMP-1, and Runx3 than the corresponding +vFcγR condition, whereas the magnitude and direction of other markers, including PD-1, NKp46, NKp44 and NKp30 varied by cluster (Fig. 6C). Because the heatmaps show z-score-normalized values, these differences reflect relative marker patterns rather than absolute protein abundance. Trajectory analysis further placed cells from the -vFcγR condition at somewhat later pseudotime values, where CD158, mTOR, T-bet, EOMES, FcεR1, and BLIMP-1 tended to increase, while NKp30 was most prominent earlier in the trajectory (Fig. 6D). Although the overlapping trajectory distributions and absence of statistical annotations preclude a claim of discrete lineage separation, the combined changes in cluster abundance and marker state indicate that HCMV vFcγRs restrain IgG-driven NK cell differentiation. In their absence, NK cells preferentially accumulate in proliferative and more differentiated effector-like states.

**Fig. 6:**
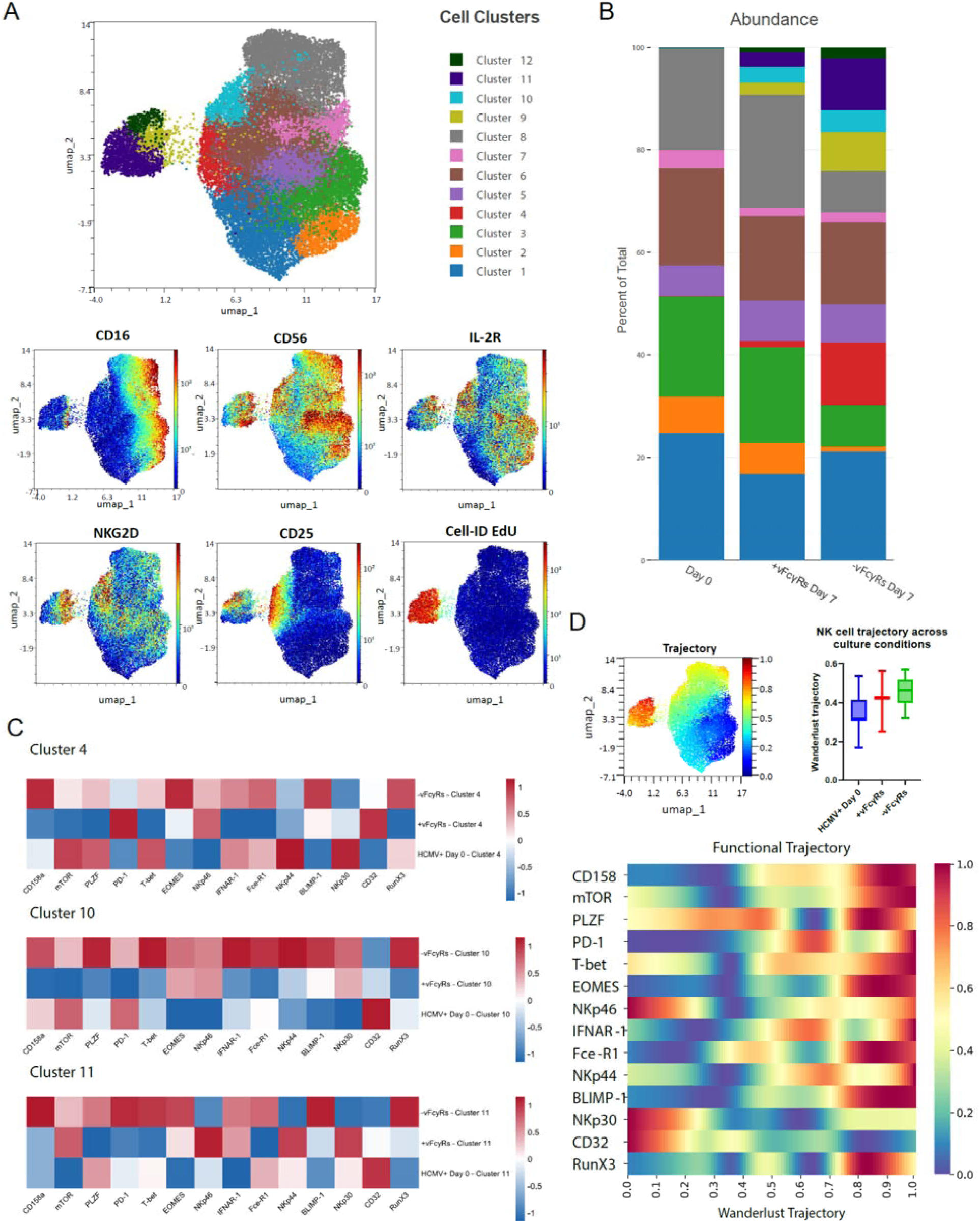
vFcγRs modulate adaptive NK cell differentiation and phenotypic imprinting. Peripheral blood mononuclear cells (PBMCs) from eight HCMV-seropositive donors were analyzed in two independent experiments with distinct donors. For downstream analysis, only donors with at least 5,000 NK cells were included, resulting in n=8 for the -vFcγRs condition and n=3 for the +vFcγRs condition. Samples were pooled for downstream analyses to ensure sufficient cell numbers and adequate representation of NK cell populations. Cells were cultured for 7 days on immobilized virions (1 x 106 PFU/ml) either containing (AD169) or lacking (AD169ΔΔΔ) or directly analyzed at 0 Days. **(A)** UMAP visualization of NK cell subsets from HCMV+ Day 0 samples (n = 8) and Day 7 cultures with virions lacking (−vFcγRs; n = 8) or containing (+vFcγRs; n = 3) vFcγRs. Samples were randomly and equally subsampled, and a single UMAP embedding was generated jointly across all HCMV+ samples. FlowSOM clustering was performed to identify distinct cell populations. **(B)** Stacked bar chart illustrating changes in NK cell cluster composition between HCMV+ Day 0 samples and Day 7 cultures following exposure to virions lacking (−vFcγRs) or expressing (+vFcγRs) viral Fcγ receptors. Cluster abundance is expressed as the percentage of total clustered cells within each experimental condition. **(C)** Heatmaps showing z-score-normalized expression of functional NK cell markers across clusters expanded at Day 7 in cultures exposed to virions lacking viral Fcγ receptors (−vFcγRs). **(D)** Wanderlust trajectory analysis was performed to investigate potential transitions between cellular states and changes in functional marker expression along the inferred trajectory, using the same markers as for dimensionality reduction. Cluster 3 from HCMV+ Day 0 samples was defined as the starting population. The inferred pseudotime trajectory is overlaid on the UMAP and color-coded according to pseudotime progression. Cultures with virions lacking viral FcγRs (−vFcγRs) project to later pseudotime states compared with cultures expressing viral FcγRs (+vFcγRs). Normalized expression of functional NK cell markers along the Wanderlust trajectory is also shown.

## Discussion

Adaptive NK cells represent specialized NK cell populations that emerges during HCMV infection and exhibits enhanced Fcγ-dependent effector function together with memory-like features (*3, 8, 9, 30, 31*). Here, we identify HCMV-encoded Fcγ-binding proteins (viral Fcγ receptors, vFcγRs) as regulators of IgG antibody-driven adaptive NK cell responses and differentiation. Three observations support this conclusion. First, the CD16a-activating capacity of HCMV-specific IgG was associated with the abundance of NKG2C+ adaptive NK cells in HCMV-seropositive individuals. Second, CD16a engagement was sufficient to preferentially expand NKG2C+ NK cells from seropositive donors. Third, gp34 and gp68 limited IgG-dependent NK cell expansion and differentiation. Expansion in the absence of vFcγRs generated more NKG2C+CD16a+ cells, greater antibody-triggered degranulation, and a shift toward proliferative and differentiated effector-like phenotypic states. These findings extend the function of vFcγRs from acute interference with FcγR activation to control of the magnitude and quality of an adaptive NK cell response.

The human cohort data suggest that the functional quality of the antibody response, rather than HCMV-specific IgG quantity alone, is associated with adaptive NK cell abundance. Total HCMV-specific IgG titers did not correlate with NKG2C+ NK cell frequency, as did neither EBV-specific CD16a activation nor HCMV-specific CD64 activation. Although the cross-sectional cohort is modest and cannot establish causality, the culture experiments provide functional support: immobilized IgG drove preferential expansion of NKG2C+ cells, soluble CD16a blockade suppressed that response, and immobilized anti-CD16a antibody was itself stimulatory. Together with previous evidence that antibody signaling promotes expansion of memory-like NK cells (*13, 14*), these results place CD16a engagement as an antibody-dependent amplification step in adaptive NK cell responses and link it mechanistically to HCMV as a trigger for adaptive NK cells (*3*) and humoral immune memory.

The experiments also connect this host amplification pathway to a defined viral immune-evasion mechanism. Virions lacking vFcγRs supported greater IgG-dependent NKG2C+ NK cell expansion than vFcγR-proficient virions. Experiments with infected cells expressing gp34 or gp68 individually, ectopic membrane expression, and soluble ectodomains further linked the suppressive activity to gp34 and gp68. The greater effect of gp68 in the infected-cell system is consistent with its ability to interfere with CD16a access to immune complexes, whereas the failure of an Fc-binding–deficient gp34 mutant to suppress expansion demonstrates that IgG-Fc engagement is required (*15, 17, 32*). Because soluble ectodomains were sufficient, internalization of antibody complexes is not necessary for suppression in this assay. The convergent results support a model in which vFcγRs reduce the availability or configuration of Fcγ for productive CD16a engagement.

The downstream consequence was not limited to cell number. NK cells that were expanded with opsonized vFcγR-deficient virions contained a larger NKG2C+CD16a+ population and showed greater degranulation after restimulation with immobilized IgG or opsonized infected cells. Mass cytometry provided a complementary view of phenotypic remodelling. In the absence of vFcγRs, proliferative CD25-high/EdU-high clusters accumulated, several clusters showed relative increases in transcription factors and receptors associated with NK cell maturation, and cells were shifted toward later pseudotime values. The overlapping trajectories and z-score-normalized marker values argue against interpreting these data as a discrete lineage transition. Instead, the flow- and mass-cytometry data together indicate that vFcγRs reshape both the abundance and state of the antibody-responsive NK cell compartment.

This antibody-dependent mechanism complements established pathways of HCMV-driven adaptive NK cell biology. Recognition of UL40-derived signal peptides presented by HLA-E, inflammatory cytokines, and accessory-cell interactions contribute to NKG2C+ NK cell expansion and persistence (*10, 30, 33, 34*), whereas stable epigenetic remodeling supports altered signaling and effector function (*2, 12, 14*). Our data suggest that HCMV-specific IgG and CD16a provide an additional amplification axis whose strength is tuned by gp34 and gp68. In this model, HCMV simultaneously supplies essential signals that favor adaptive NK cell responses but encodes also vFcγRs that limit their antibody-driven expansion and maturation. Such opposing viral activities may help explain the marked interindividual variation in adaptive NK cell frequency after HCMV exposure.

Several limitations define the scope of these conclusions. The cohort analysis is cross-sectional, and the expansion assay models antibody stimulation *ex vivo* rather than adaptive NK cell development during infection *in vivo*. NKG2C was used as the operational marker of adaptive NK cells; this definition does not capture NKG2C-negative adaptive states and excludes individuals with homozygous KLRC2 deletion who also exhibit adaptive NK cell populations (*30*).

By linking vFcγRs to the composition and responsiveness of an adaptive immune-cell population, this study broadens the concept of FcγR immune evasion beyond transient inhibition of antibody effector function. The data establish gp34 and gp68 as brakes on the IgG/CD16a-driven adaptive NK cell axis and provide a framework for determining whether vFcγR targeting can strengthen antibody-based immunity or CMV-vector platforms.

## Materials and Methods

### Human subjects and blood samples

PBMCs were obtained with informed consent from healthy volunteers or from de-identified buffy coats supplied by University Hospital Freiburg.

Genomic DNA was isolated from 200 µL blood with a TanBead extractor (Taiwan Advanced Nanotech Inc.) and eluted in 50 µL nuclease-free H₂O. KLRC2wt/del variants were determined by touchdown PCR (*35*), and FCGR3A-V/F158 (rs396991) variants with a QuantStudio 5 real-time PCR system (Applied Biosystems, Darmstadt, Germany), TaqMan SNP Genotyping Assays, and TaqMan Genotyping Master Mix. The study followed federal guidelines, Albert-Ludwigs-University Freiburg ethics regulations (approval 474/18), and the 1975 Declaration of Helsinki. Donor plasma serostatus was determined with virus-specific ELISA kits, and PBMCs were selected by availability (Supplementary Tables 1 and 2). The HCMV-positive cohort (n=59) was 52.5% female (median age 27.5 years) and 47.5% male (median age 27 years); the HCMV-negative cohort (n=16) was 56% female (median age 26 years) and 44% male (median age 24 years).

**Table 1.** Clinical Information/history of serum donors. Serum was collected from healthy HCMV seropositive and seronegative individuals. na= information not available, f= female, m= male

| Donor no. | CMV status | Sex | Age at time of sampling | CMV IgG [AU/m] | %NKG2C+ NK cells at time of sampling | EBV [S/co] | Genotype CD16a | Genotye NKG2C |
|---|---|---|---|---|---|---|---|---|
| 5 | positive | m | 29 | 81,70 | 20,8 | 21,46 | F/V | wt/del |
| 6 | positive | f | 30 | 97,50 | 77,7 | 19,78 | F/V | wt/wt |
| 10 | positive | f | 50 | 22,90 | 6,02 | 2,93 | F/V | del/del |
| 14 | positive | f | 26 | 59,60 | 72 | 15,16 | V/V | wt/wt |
| 15 | positive | m | 35 | 70,70 | 18,1 | 16,54 | F/V | wt/wt |
| 28 | positive | m | 27 | 109,00 | 33,3 | 11,39 | F/F | wt/wt |
| 30 | positive | m | 49 | 131,00 | 45,7 | 22,39 | V/V | wt/wt |
| 39 | positive | f | 31 | 75,80 | 23,5 | 68,28 | F/F | wt/del |
| 43 | positive | f | 25 | 51,70 | 20,2 | 20,43 | n.a. | n.a. |
| 44 | positive | f | 25 | 127,40 | 10,6 | n.a. | F/F | wt/wt |
| 46 | positive | f | 25 | 199,80 | 48,6 | 18,96 | F/V | wt/wt |
| 47 | positive | m | 27 | 190,60 | 8,22 | n.a. | V/V | wt/wt |
| 48 | positive | f | 55 | 69,90 | 11 | n.a. | F/F | wt/wt |
| 50 | positive | f | 28 | 142,20 | 2,85 | 0,04 | F/V | wt/del |
| 53 | positive | f | 55 | 212,00 | 15,8 | 5,8 | F/F | wt/wt |
| 56 | positive | f | 26 | 79,20 | 16,3 | 40,6 | F/V | del/del |
| 57 | positive | f | 27 | 122,00 | 18,6 | n.a. | F/F | wt/wt |
| 58 | positive | m | 25 | 145,00 | 27,6 | n.a. | F/V | wt/wt |
| 61 | positive | f | 25 | 113,60 | 18,1 | 28,24 | F/F | wt/wt |
| 62 | positive | f | 25 | 121,60 | 4,02 | 30,94 | F/V | del/del |
| 66 | positive | m | 25 | 76,8 | 23,7 | 8,4 | F/F | wt/wt |
| 76 | positive | f | 22 | n.a. | 42,6 | n.a. | F/V | wt/wt |
| 77 | positive | f | 27 | 51,2 | 27,1 | 18,44 | F/F | wt/wt |
| 78 | positive | f | 27 | 19,70 | 6,58 | 21,54 | F/V | wt/wt |
| 94 | positive | n.a. | n.a. | n.a. | 35,1 | n.a. | F/V | wt/wt |
| 96 | positive | n.a. | n.a. | n.a. | 15,3 | n.a. | F/V | wt/wt |
| 97 | positive | n.a. | n.a. | n.a. | 41,4 | n.a. | F/F | wt/wt |
| 100 | positive | n.a. | n.a. | n.a. | 17,8 | n.a. | F/V | wt/wt |
| 101 | positive | m | 46 | 180 | 10,64 | 19,63 | F/V | wt/wt |
| 33 | negative | f | 28 | <5,00 | 10,4 | <5,00 | n.a. | n.a. |
| 36 | negative | m | 23 | <5,00 | 27 | 0,05 | V/V | wt/wt |
| 45 | negative | f | 25 | <5,00 | 4,89 | 18,96 | F/F | wt/wt |
| 52 | negative | m | 26 | <5,00 | 14 | 0,05 | F/V | wt/wt |
| 55 | negative | m | 24 | <5,00 | 9,78 | 16,12 | F/F | wt/wt |
| 59 | negative | m | 30 | <5,00 | 7,24 | 28,82 | V/V | wt/wt |
| 63 | negative | m | 25 | <5,00 | 6,73 | 0,05 | F/F | wt/wt |
| 65 | negative | f | 28 | <5,00 | n.a. | 0,05 | F/V | wt/wt |
| 67 | negative | f | 26 | <5,00 | n.a. | 0,05 | F/F | wt/del |
| 69 | negative | f | 25 | <5,00 | n.a. | 22,48 | F/V | wt/del |
| 70 | negative | f | 27 | <5,00 | n.a. | 5,84 | F/V | wt/del |
| 71 | negative | f | 28 | <5,00 | n.a. | 0,05 | F/V | wt/del |
| 72 | negative | f | 25 | <5,00 | n.a. | 0,05 | F/V | wt/wt |
| 73 | negative | f | 24 | <5,00 | n.a. | 0,05 | V/V | wt/del |
| 74 | negative | m | 23 | <5,00 | n.a. | 0,05 | F/V | na |
| 75 | negative | m | 23 | <5,00 | n.a. | 44,61 | V/V | wt/wt |

**Table 2.** Selection of donor samples per experiment. The donor material was selected for individual experiments according to availability or potential to expand adaptive NK *in vitro*. The donor material is unmatched for gender or age.

|  | **1B** | **1C-H** | **2 C-D** | **4A** | **4B** | **4C-D** | **5B-D** | **5E** | **5F-G** | **6A-D** | **S2E** | **S4A** | **S4B** | **S4C** | **S5A** | **S5B** | **S5A** |
|---|---|---|---|---|---|---|---|---|---|---|---|---|---|---|---|---|---|
| 5 | x | x | x |  |  |  |  |  |  |  | x |  | x | x | x |  |  |
| 6 | x | x | x |  |  |  |  |  |  |  | x |  | x | x | x |  |  |
| 10 | x |  | x |  |  |  |  |  |  |  | x |  |  |  |  |  |  |
| 14 | x | x | x | x |  | x | x | x | x | x | x |  |  |  |  | x | x |
| 15 | x | x | x | x |  | x |  |  | x | x | x | x | x | x |  |  | x |
| 28 | x | x | x |  |  |  |  |  |  |  | x |  |  |  |  |  |  |
| 30 | x | x | x | x | x | x | x |  | x | x | x |  | x | x | x |  | x |
| 39 | x | x | x | x |  | x |  |  | x |  | x |  |  |  |  |  | x |
| 43 | x |  | x |  |  |  |  |  |  |  | x |  |  |  |  |  |  |
| 44 | x | x | x | x |  |  |  |  |  |  | x |  | x | x | x |  |  |
| 45 | x |  |  | x |  |  |  |  |  |  |  |  |  | x |  |  |  |
| 46 | x | x | x |  |  |  |  |  |  | x | x |  | x |  | x |  |  |
| 47 | x | x | x |  |  |  |  |  |  |  |  |  |  | x | x |  |  |
| 48 | x | x | x |  |  |  |  |  |  |  |  |  | x |  | x |  |  |
| 50 | x | x | x |  |  |  |  |  |  |  |  |  |  |  | x |  |  |
| 53 | x | x | x |  |  |  |  |  |  |  |  |  |  |  | x |  |  |
| 56 | x | x | x |  |  |  |  |  |  |  |  |  |  |  | x |  |  |
| 57 | x |  | x |  |  |  |  |  |  |  |  |  |  |  | x |  |  |
| 58 | x |  | x |  |  |  |  |  |  |  |  |  |  |  | x |  |  |
| 61 | x | x | x | x |  | x |  |  | x | x |  |  | x | x | x |  | x |
| 62 | x | x |  |  |  |  |  |  |  |  |  |  |  |  |  |  |  |
| 66 | x | x |  |  |  |  |  |  |  |  |  |  | x | x |  |  |  |
| 76 | x | x |  |  |  |  |  |  |  |  |  |  |  |  |  |  |  |
| 77 | x |  |  | x |  | x |  |  |  |  |  |  |  |  |  |  | x |
| 78 | x | x |  |  |  |  |  |  |  |  |  |  |  |  |  |  |  |
| 85 |  |  |  |  |  |  |  |  | x |  |  |  |  |  |  |  |  |
| 94 | x |  |  | x |  | x | x |  | x | x |  |  | x | x |  |  | x |
| 96 |  |  |  |  |  |  | x |  | x | x |  |  |  |  |  |  |  |
| 97 | x |  |  | x |  | x | x |  | x | x |  |  | x | x |  |  | x |
| 100 |  |  |  |  |  |  |  |  |  |  |  |  | x | x |  |  |  |
| 101 | x | x |  |  |  |  |  |  |  |  |  |  |  |  |  |  |  |
| 33 | x |  | x |  |  |  |  |  |  | x |  |  |  |  |  |  |  |
| 36 | x |  | x |  |  |  |  |  |  | x |  |  |  |  |  |  |  |
| 52 | x | x |  |  |  |  |  |  |  |  |  |  |  |  |  |  |  |
| 55 | x | x |  |  |  |  |  |  |  | x |  |  |  |  |  |  |  |
| 59 | x | x |  |  |  |  |  |  |  | x |  |  |  |  |  |  |  |
| 63 | x |  | x |  |  |  |  |  |  | x |  |  |  |  |  |  |  |
| 65 | x |  | x |  |  |  |  |  |  |  |  |  |  |  |  |  |  |
| 67 | x |  | x |  |  |  |  |  |  | x |  |  |  |  |  |  |  |
| 69 | x | x | x |  |  |  |  |  |  |  |  |  |  |  |  |  |  |
| 70 | x |  | x |  |  |  |  |  |  |  |  |  |  |  |  |  |  |
| 71 | x |  | x |  |  |  |  |  |  |  |  |  |  |  |  |  |  |
| 72 | x |  | x |  |  |  |  |  |  |  |  |  |  |  |  |  |  |
| 73 | x |  | x |  |  |  |  |  |  |  |  |  |  |  |  |  |  |
| 74 | x |  | x |  |  |  |  |  |  |  |  |  |  |  |  |  |  |
| 75 | x | x |  |  |  |  |  |  |  |  |  |  |  |  |  |  |  |
| 76-88 | x |  |  |  |  |  |  |  |  |  |  |  |  |  |  |  |  |

### Cell Isolation and Cell culture

293T (human embryonic kidney; ATCC CRL-11269), HFF (HF-99/7), ARPE-19 (retinal pigment epithelium; ATCC CRL-2302), and MRC-5 (human lung fibroblast; ATCC CCL-171) cells were cultured in DMEM (Gibco) with 10% heat-inactivated FCS, 100 U/mL penicillin, 100 µg/mL streptomycin (Gibco), and 50 µM β-mercaptoethanol. BW5147 mouse thymoma cells were maintained at 3 × 10LJ–9 × 10LJ cells/mL in RPMI GlutaMAX (Gibco) with 10% FCS, 1× sodium pyruvate (Gibco), and 0.1 mM β-mercaptoethanol (Sigma). P3HR1 cells (provided by Valeria Kapper-Falcone, Institute of Virology, Freiburg, Germany) were cultured in RPMI with 20% FCS, 500 U/mL penicillin, 500 µg/mL streptomycin, and 0.4% 200 mM L-glutamine (Gibco). PBMCs were isolated by density-gradient centrifugation in lymphocyte-separation medium (Anprotec), used immediately or cryopreserved in FCS/10% DMSO, and maintained in NK MACS Medium (Miltenyi Biotec) with 1 µg/mL ciprofloxacin and 1 IU/mL IL-2, or in GMP Stem Cell Growth Medium (SCGM; CellGenix); both contained 10% heat-inactivated HCMV-seronegative human serum. All cells were cultured at 37°C and 5% CO₂.

### Preparation of soluble vFcγRs

293T cells were seeded at 70% confluence one day before transfection with polyethylenimine (Sigma-Aldrich) according to the supplier’s instructions. Constructs were expressed from pIRES-eGFP (Addgene). After 3 days at 37°C, supernatants were collected, stored at -80°C, and quantified for soluble vFcγRs by western blotting (*15*).

### Virus stock preparation and infection of cells

The bacterial artificial chromosome (BAC) derived from the HCMV strain AD169 was used as a parental virus, from which a triple deletion virus (termed here as ΔΔΔ) lacking genes *RL11*, *RL12* and *UL119-118* was generated previously (*34*). TB40/E strain (*28*) (kindly provided by Christian Sinzger, Ulm) was pre-cultured on ARPE-19 cells to retain epithelial tropism and pentameric complex (PC) expression according to previous observations (*36*). Virus stocks were grown by infecting MRC-5 cells with a multiplicity of infection (MOI) of 5. After 7 days of incubation, the supernatant was collected and the infected cells were scraped off the flask and washed off with 10 ml of supernatant, then added to the rest of supernatant. The cell suspension was dounced and spinned down. Resulting supernatant was added to supernatant and stored at -80°C. Virus titers were determined by titrating virus stocks onto MRC5 cells and detecting the immediate early antigen 1(IEA1) expression with a monoclonal antibody (anti HCMV I.E.A., Argene Biosoft) in fluorescence microscopy. For infection, MRC-5 or HFF cells were seeded one day before in DMEM with 10% FCS. Cells were infected by centrifugal enhancement (2x15min at 1.200g).

### NK cell proliferation assay

PBMCs were purified from healthy donor and cultivated as described. Prior cultivation, cells were stained with a proliferation dye (Celltrace, Invitrogen) with a concentration of 5–10 μM/ 1*10^6^ cells, according to manufacturer’s instructions. PBMCS were then incubated with opsonized IgG in different set ups as listed and analyzed after 7 days via flow cytometry: <u>Immobilized human IgG:</u> human or humanized IgG was added with a concentration of 50-0µg/ml to a 96well Maxisop ELISA plate and was incubated for 1h at RT.

#### IgG-opsonized HCMV Virions

HCMV virions were opsonized via incubation on a Maxiosorp 96 well ELISA Plate at 37°C for 1h. HIG was added to the well with a concentration of 50µg/ml and incubated for 1h at 37°C

#### IgG-opsonized transfected Hek293T

constructs of gp34, gp68 and CD20 were expressed on Hek293T cells by transfection opsonized with antiCD20 10µg/ml antibody (rituximab)

#### IgG-opsonized HCMV infected cells

MRC-5 cells were infected as described earlier and opsonized with 50µg/ml HIG

### FcγR activation assay

MRC-5 cells were seeded in 96 well plates before infected using TB40/E at MOI = 4 for 72h. For EBV, P3HR1 cells (1*10^6^ cells/mL) constitutively expressing EBV antigens were seeded in a 96 well format. Graded amounts of antibody or serum were incubated with target cells for 1h, followed by replacement of medium and co-culture with FcγR BW5147-reporter cells for 16h (E:T ratio: 20:1). All incubation steps were performed at 37°C in a 5% CO_2_ atmosphere. Reporter cell mIL-2 secretion was quantified by subsequent anti mouse IL-2 sandwich ELISA as described previously (*37*).

### Flow Cytometry

Cell suspensions were stained with combinations of the following conjugated antibodies according to manufacturing guidelines: CD159c (NKG2C)/PE REAinfinity (1:100), CD56 APC (1:200), CD159a (NKG2A) Biotin, REAfinity™ (1:100; all Miltenyi Biotec), CD3 PerCP/Cyanine5.5 (1:50), CD16a B421/BV711 (1:100), CD107a BV711 (1:50). Streptavidin BV421 (1:100; all BioLegend), anti-FcεR, γ subunit-FITC (1:50; Merck), CD57 eBioscience Pe-Cyanine 7 (1:100; Invitrogen). Proliferation was tracked by CellTrace CFSE or Violet Proliferation stain (dye concentration of 5–10 μM/ 1*10^6^ Cells; Invitrogen). Dead cells were excluded using LIVE/DEAD Fixable Red stain (Thermo Fisher) or a Zombie NIR™ Fixable Viability kit (BioLegend). Cytometry was performed using a LSR Fortessa instrument (BD Bioscience) and data was analyzed via FACS-Diva (BD) and FlowJo (TreeStar Inc.) software. Dimension reduction algorithms “Uniform Manifold Approximation and Projection” and “t-distributed stochastic neighbor embedding”(*38*) were used to visualize acquired data. FlowSOM population clustering was applied to categorize and quantify populations (*29*).

### NK cell degranulation assay

PBMCs were purified from healthy donor blood by centrifugation via Lymphoprep Medium according to the supplier instructions (Anprotec). HCMV-infected MRC-5 or HFF cells (MOI = 3, 72hpi) were incubated with titrated amounts of Cytotect® (Biotest, Dreieich, Germany) at 37°C for 1 hr. After washing twice with medium, 5 × 10^5^ PBMCs were incubated on opsonized infected cells for 6 hrs (100 µl per well in RPMI/10% FCS) in the presence of αCD107a-, αCD56-BV650, and Golgi-Plug/Golgi-Stop (according to supplier, BD). After incubation, PBMCs were harvested, washed in staining buffer (PBS/3% FCS), and incubated with αCD3-FITC 30 min at 4°C.

### Mass Cytometry (CyTOF)

Mass Cytometry was performed by the CyTOF facility at the university medical center Freiburg as described (*39*). In brief, samples were first barcoded using a palladium-based barcoding strategy according to the manufacturer’s instructions to allow multiplexed sample acquisition and minimal batch variation. Following barcoding, cells were exposed to a viability assay (MM-DOTA) to exclude dead cells. To assess proliferation, cells were stained for Cell-IDU incorporation. Subsequently, cells were stained with a panel of metal-conjugated antibodies targeting extracellular (surface) markers. After surface staining, cells were fixed and permeabilized using (Foxp3/Transcription Factor Staining Buffer Set), followed by staining for intracellular markers where indicated. Cells were then incubated with an iridium-based DNA intercalator for identification of single, nucleated cells. Complete CyTOF NK-Cell Panel is provided in the supplementary material (Fig. S8). Samples were then acquired on a Helios mass cytometer (Fluidigm) Data were normalized using the CyTOF software and analyzed using FlowJo and Omiq Software. Gaussian Parameter Data Cleanup was performed followed by debarcoding and FCS files were imported into OMIQ (Dotmatics; https://www.omiq.ai) for downstream analysis. Data were arcsinh-transformed using a cofactor of 5. Following manual gating, viable singlet CD3-CD14-CD19-CD56+ NK cells were identified for further analysis. For dimensionality reduction, samples were randomly and equally subsampled, and a single UMAP embedding was generated jointly across all HCMV+ samples. FlowSOM clustering was performed to identify distinct cell populations. Wanderlust Trajectory analysis was subsequently used to investigate potential transitions between cellular states and changes in functional marker expression along the inferred trajectory. Trajectory analysis was performed using the same markers included in the dimensionality reduction. Cluster 3 from HCMV+ Day 0 samples was defined as the starting population for trajectory inference.

### Large Language Models

ChatGPT (OpenAI) was used for language editing and condensation of the manuscript. The authors reviewed and revised all generated text and take full responsibility for the content.

## Data availability

All data needed to evaluate the conclusions of this study are available in the paper and/or the Supplementary Materials. Additional materials are available from the corresponding author upon reasonable request.

## Acknowledgments

This work was funded by the German Research Foundation (through DFG KO6815/1-1, HE2526/9-1 and HE2526/9-2) awarded to PK and HH. PK has received the Renate-And-Hans-Schleussner Award from Biotest AG (Frankfurt, Germany).

## Author contributions

Conceived and designed the experiments: MH, PK, HH Performed the experiments: MH, NS, SH, MS, MU, HV Analyzed the data: MH, NS, PK, MU, HV, POM, GP Writing and original draft preparation: MH, PK Review and editing: HH, EPS, MLB, BB, PK

## Competing interests

PK has received the Hans and Renate Schleussner Award by Biotest GmbH & Co. KGAa and has received Cytotect® for research purposes. None of these activities have impacted this work beyond financial and material support. The remaining authors declare no competing interests.

## Supplementary Information

**Fig. S1.**
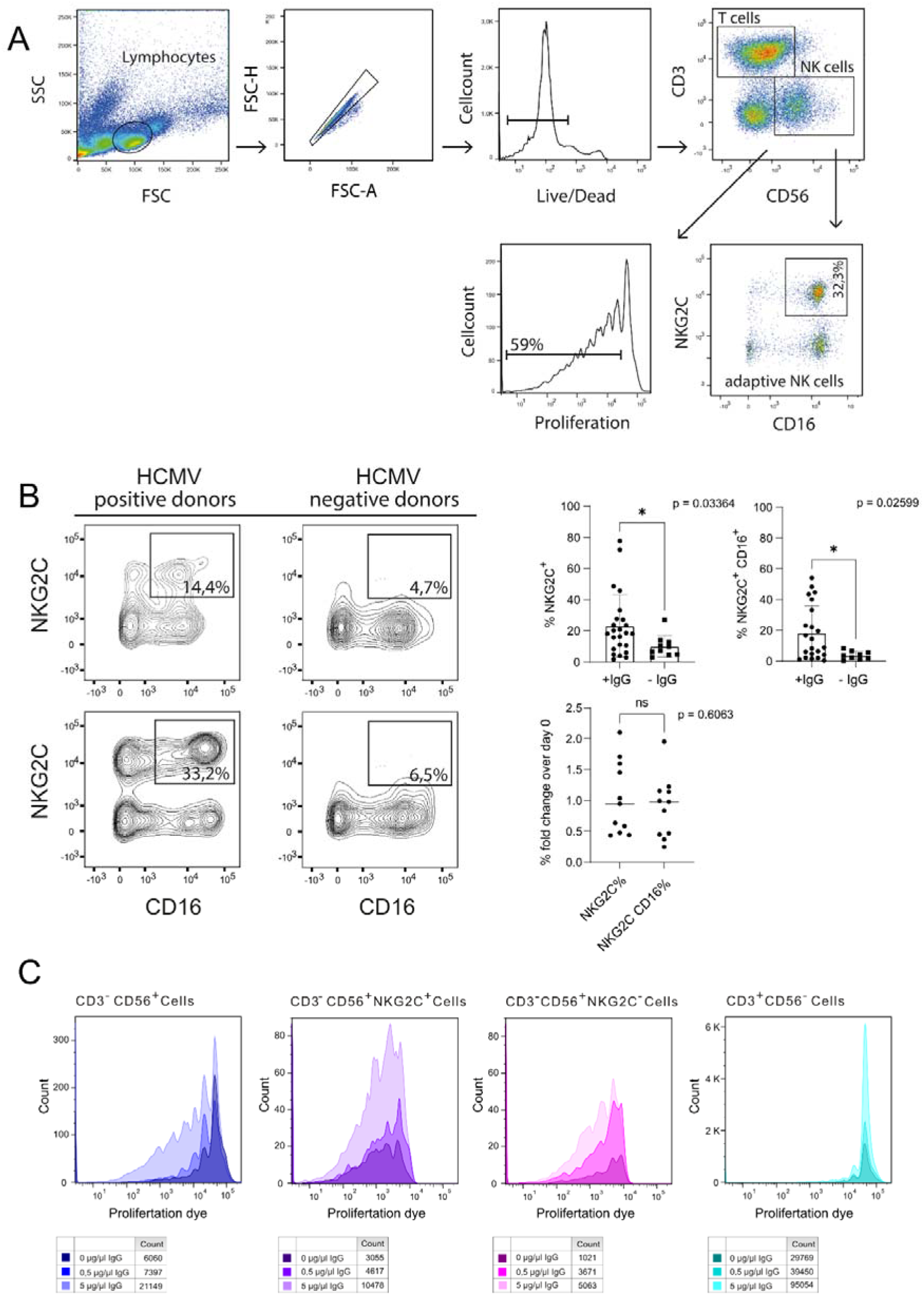
Gating strategy and characterization of adaptive NK cells within the cohort. **(A)** Representative gating strategy measuring cell proliferation and adaptive NK cell marker expression is shown. NK cells expressing a high level of NKG2C alone or with CD16a are identified as adaptive NK cells. Changes in the percentage of NKG2C positive NK cells were used as measurement to detect adaptive NK cell expansion. Further a CFSE pre-staining was performed to track proliferation. Proliferation was assessed by either MFI or determined as percentage of divided cells. **(B)** (right) exemplary NK cell populations from HCMV positive and negative healthy donors showing an increased proportion of NK cells with high expression levels of NKG2C and CD16a in positive individuals. (left) Among a HCMV seropositive and negative cohort, percentage of NKG2C^+^ NK cells (up left) and CD16a^+^ NKG2C^+^ NK cells (up right) is compared. Fold change increase of IgG mediated expansion for 7 days over day 0 was compared of NKG2C^+^ and NKG2C^+^CD16a^+^ to determine expansion potential (bottom). To test statistical significance, Mann Whitney test (two tailed) as used **(C)** Exemplary depiction of the measurement of IgG induced proliferation. PBMCs were pre-stained with Cell Trace™ violet proliferation dye and cultured in a sterile ELISA plate coated with titrated amounts of anti CD20 IgG (rituximab). After 7 days of incubation, proliferation of NK cells (CD3^-^ CD56^+^) compared to T cells (CD3^+^ CD56^-^) was detected via flow cytometry.

**Fig. S2.**
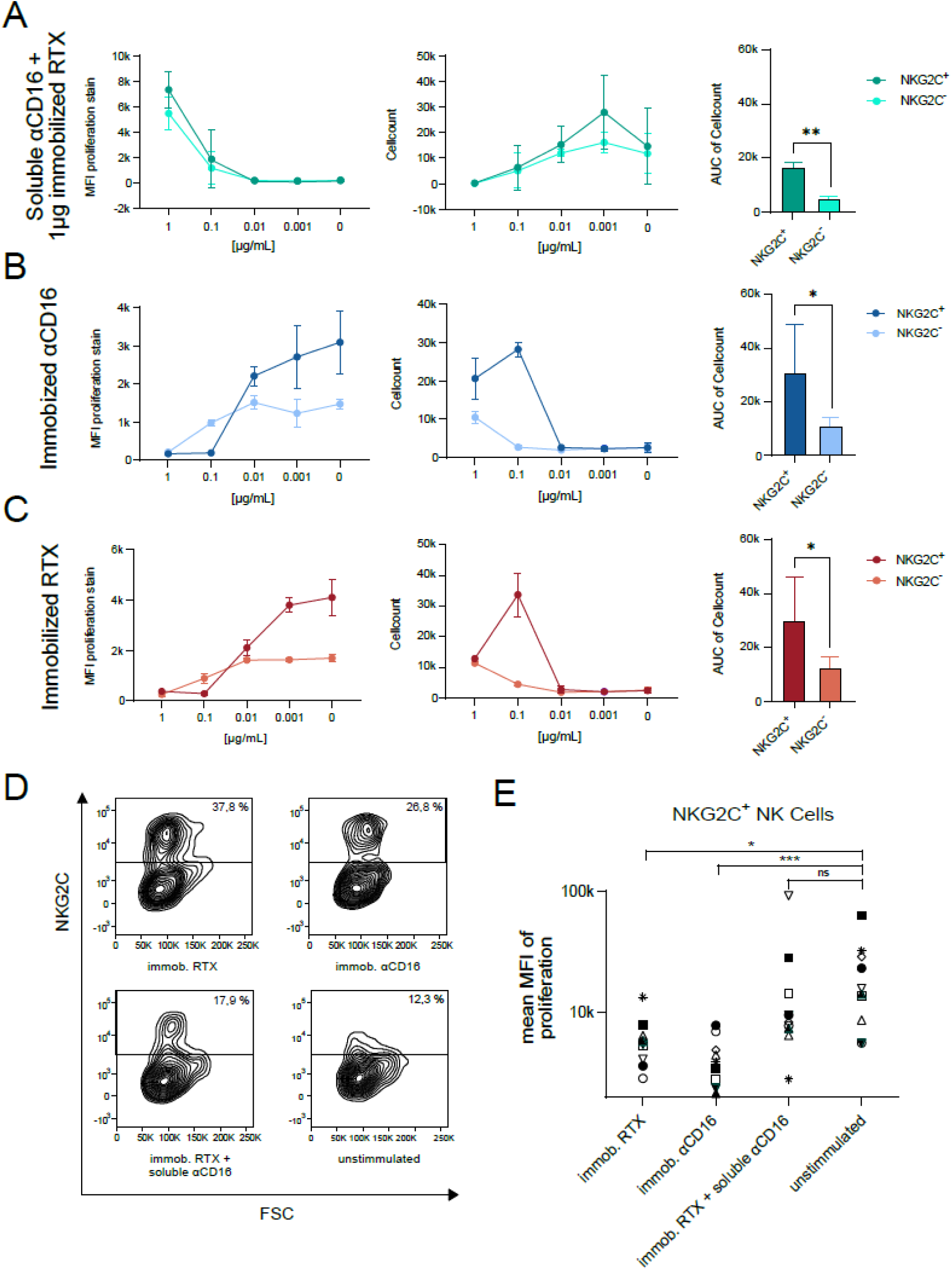
Blocking of CD16a inhibits adaptive NK cell expansion. PBMCs of one representative donor were probed for proliferation in the presence of **(A)** 1µg of immobilized human IgG1 (rituximab) in combination with 1µg of soluble anti-CD16a IgG **(B)** immobilized anti-CD16a IgG (3G8) **(C)** or 1µg of immobilized human IgG1 (rituximab) alone. After 7 days of culture, proliferation and total cell count of NKG2C^+^ or NKG2C^-^ NK cells were measured. Area under curve (AUC) values summarizing the cell counts across the titration are shown as bar graphs. Error bars = SD, two independent experiments. **(D)** Cytometry density plots showing NKG2C expression by NK cells (CD3^-^/CD56^+^) of one exemplary donor at 1µg of immobilized rituximab, 1µg of immobilized 3G8 or 1µg of immobilized rituximab in combination with 1µg of 3G8. Unstimulated cells are shown for reference. **(E)** Percentage of NKG2C^+^ NK cells (left) and proliferation (CellTrace™ MFI, right) of an HCMV sero-positive cohort (n=11) tested as above **(A-C)**.unpaired t-Test, one tailed **(E)** Wilcoxon matched pairs ranked sign test, two-tailed. ***=p<0.0002**=p<0.0021, *=p<0.0332,

**Fig. S3.**
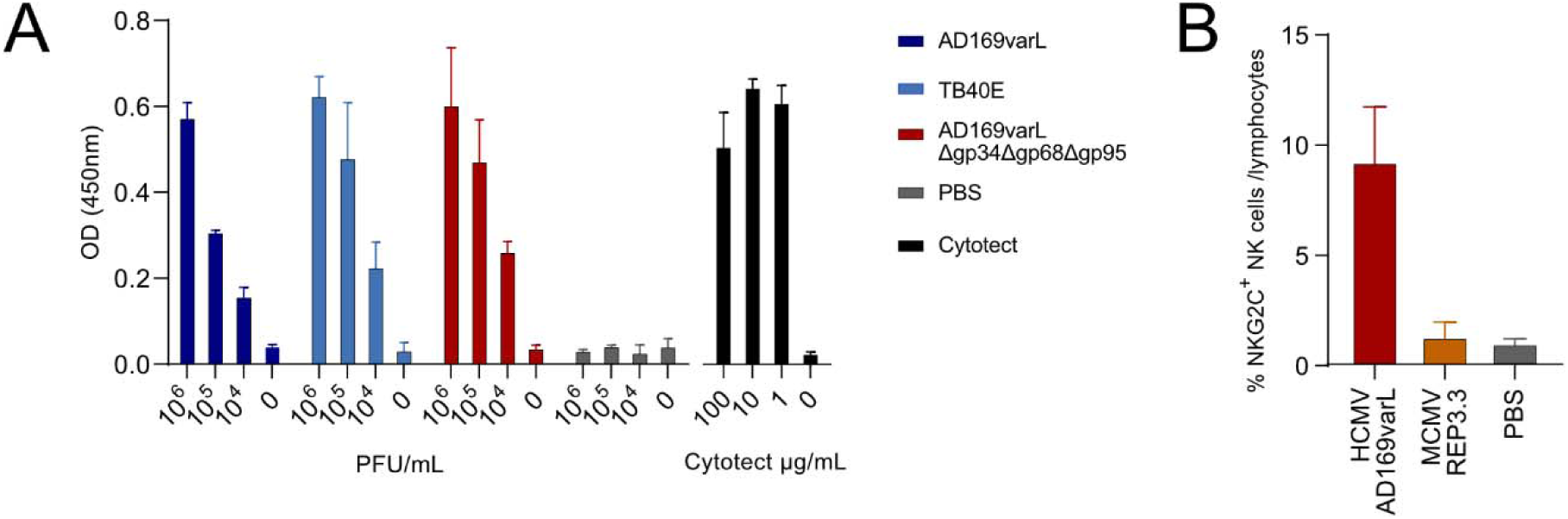
HIG opsonized HCMV virions specifically and comparable opsonization between HCMV variants required tuning of HIG usage. **(A)** Virions of HCMV variants opsonized using HIG (Cytotect^®^ 1μg), or HIG alone, were immobilized in a dilution series. The amount of immobilized IgG was then detected by human IgG specific ELISA (HRP). Optical density was measured at a wavelength of 450 nm. **(B)** PBMCs were cultured for 7 days in the presence of HIG (1μg) on immobilized virions of HCMV AD169 or MCMV strain REP3.3 (1.15*106 PFU) or PBS, followed by cytometry analysis of NKG2C+ NK cell expansion (CD3^-^/CD56^+^). Two independent experiments in technical duplicates. Error bars = SEM

**Fig. S4.**
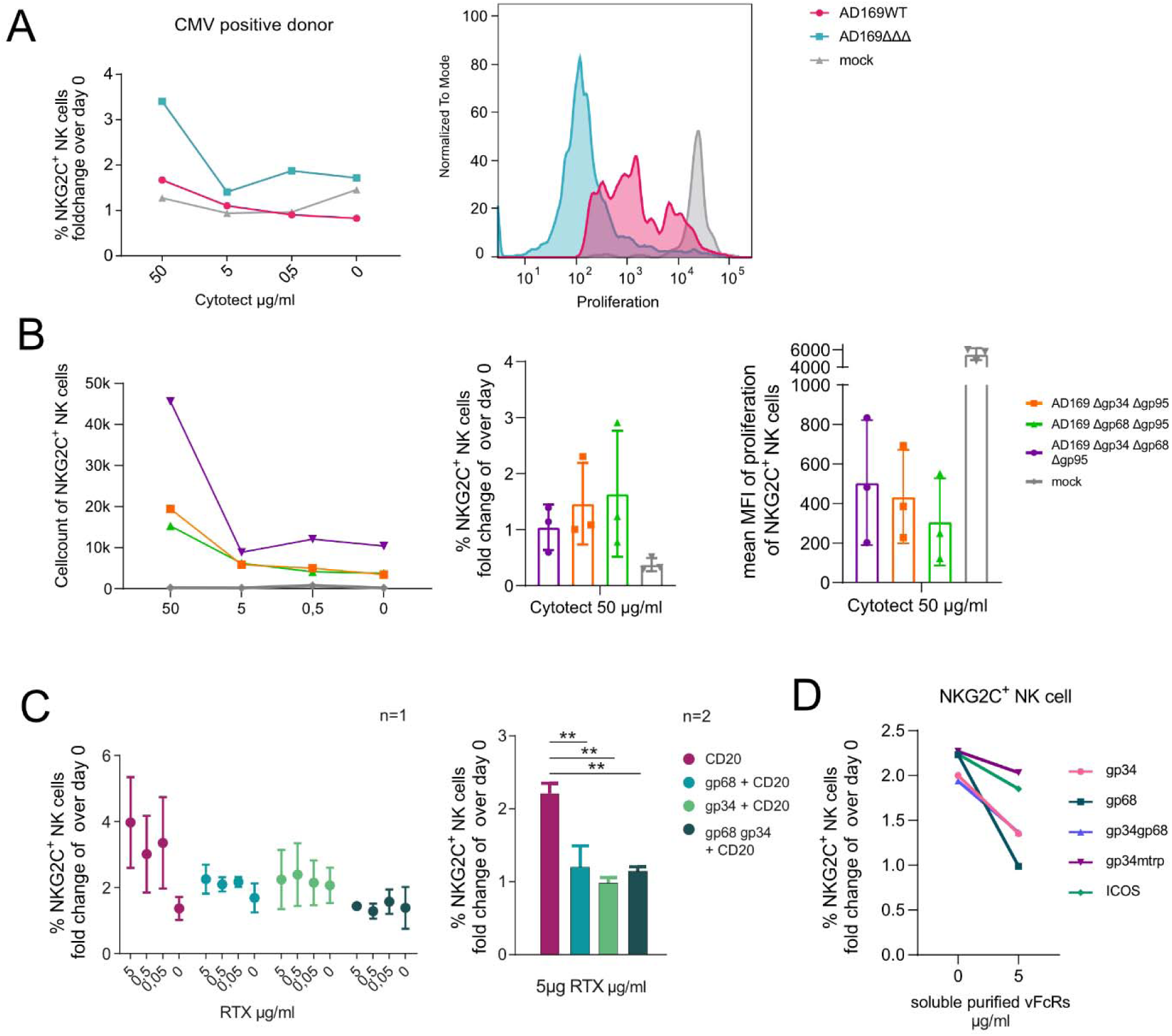
Soluble and membrane resident forms of gp68 and gp34 limit adaptive NK cell expansion. **(A)** HFF cells were infected with HCMV AD169 or AD169ΔΔΔ or mock infected (MOI = 5). Incubation with graded amounts of HIG (Cytotect^®^) was followed by incubation with PBMCs from one donor prestained with celltrace violet proliferation dye. After 7 days of incubation, cells were analyzed via flow cytometry. Depicted are relative frequencies of NKG2C^+^ NK cells related to total lymphocyte population (left) and the MFI of the proliferation stain of the NKG2C^+^ NK cell population with 0.1μg Cytotect^®^ (right). **(B)** HFFs were infected with AD169Δgp34Δgp95 (ΔRL11/ΔRL12) or AD169Δgp68Δgp95 (ΔUL119-118/ΔRL12) or AD169ΔΔΔ deletion variants and treated and analyzed for NKG2C^+^ cell expansion using PBMCs from a representative donor. Graded amounts of HIG were used. Shown are4 cell counts of NKG2C^+^ NK cells measured with equal sample volume and time period (left). Mean of the percentage of NKG2C^+^ NK cells (normalized to day 0) (center) and MFI of proliferation are shown (n=3) (right). **(C)** 293T cells were transfected with a CD20 alone or co-expressed with CD99 as a non-Fcγ binding control molecule. HCMV vFcγR gp68, gp34 or both were co-expressed prom a p_IRES-eGFP expression vector as indicated. Cells were incubated with 1μg humanized anti-CD20 monoclonal IgG (rituximab, RTX). PBMCs of HCMV positive donors were added and co-cultured for 7 days. NKG2C^+^ adaptive NK cell frequency normalized to total lymphocytes are showed as fold increase over day 0. Two independent experiments of the same donor (left). Bar graphs compare values at 5µg/ml immobilized RTX. Means from two individual HCMV positive donors. Error bars = SEM. **(D)** HER-2 antigen positive SKOV-3 cells were incubated with 1µg humanized anti-Her2 monoclonal IgG (trastuzumab, Herceptin^®^). Supernatants from soluble vFcγR producing 293T cells were given in titrated amounts. The Fcγ binding deficient point mutant of gp34 (gp34mtrp) was used as a negative control. PBMCs of an HCMV positive donor were co-cultured on opsonized cells for 7 days. Frequencies of NKG2C^+^ adaptive NK cells normalized to day 0 as reference is shown. *=p<0.05; **=p<0.01; ***=p<0.001 (1-way analysis of variance (ANOVA), Fishers LSD).

**Fig. S5.**
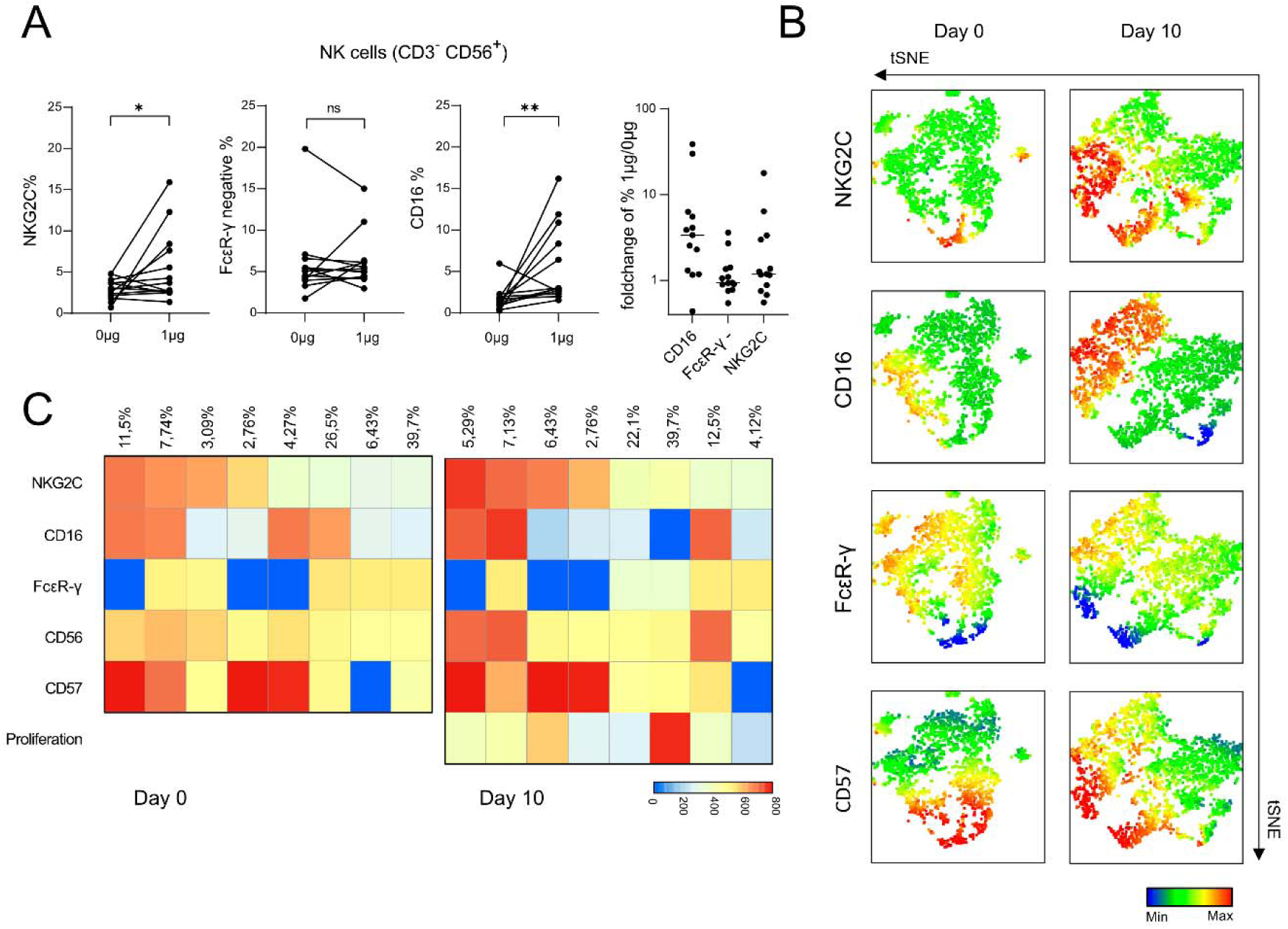
Analysis of IgG induced expansion potential in NK cell populations. **(A)** PBMCs of HCMV positive donors (n=13) were cultures with 1µg immobilised IgG for 7 days to determine the expansion potential in regard of marker expression. **(B)** NK cells of a representative donor are shown before (day 0) and after (day 7) IgG mediated expansion and visualized using t-SNE. **(C)** Pooled analyses integrating data from 8 donors as in **(A)** before and after expansion on 1µg immobilized. IgG. Before data integration, counts were set to 1.000 cells to ensure equal representation. Population demarcation was performed using FlowSOM and populations were analysed for expression of NKG2C, CD16a, FcεR γ, CD56, CD57 and proliferation. Heat map shows rates of all populations classified via FlowSOM before (d0) and after cultivation as in **(A)**. Color scale indicates MFI.

**Fig. S6.**
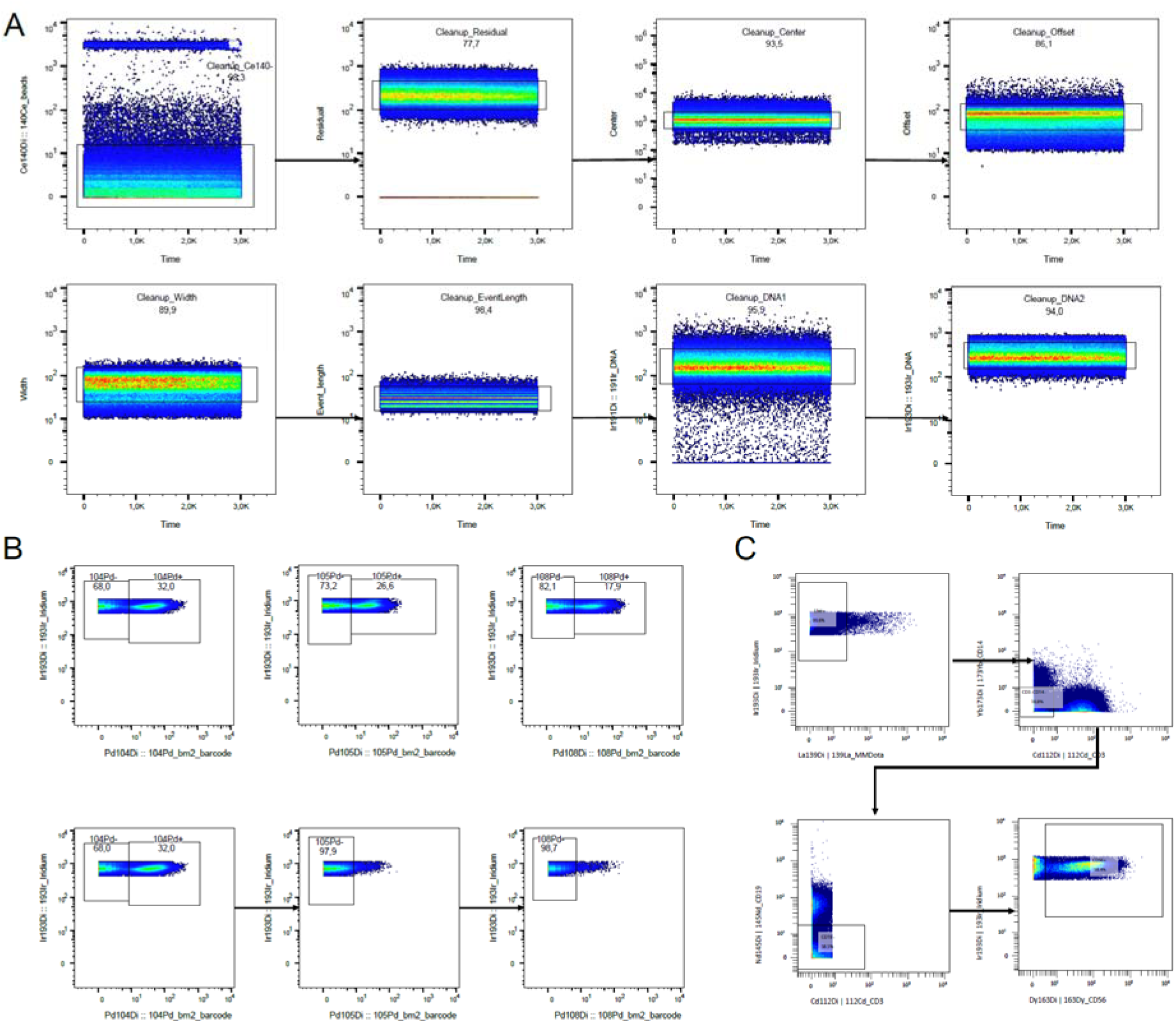
CyTOF Data Cleanup, Debarcoding and Gating Strategy. **(A)** CyTOF data were cleaned using Gaussian parameters to exclude events with abnormal acquisition profiles and reduce technical artifacts. **(B)** A combinatorial metal-barcode scheme was used to uniquely label individual samples, allowing samples to be pooled for staining and acquisition. Each sample was defined by one positive and two negative barcode gates, enabling reliable sample identification. **(C)** Representative gating strategy used to identify CD56⁺ NK cells for downstream multidimensional analysis.

**Fig. S7.**
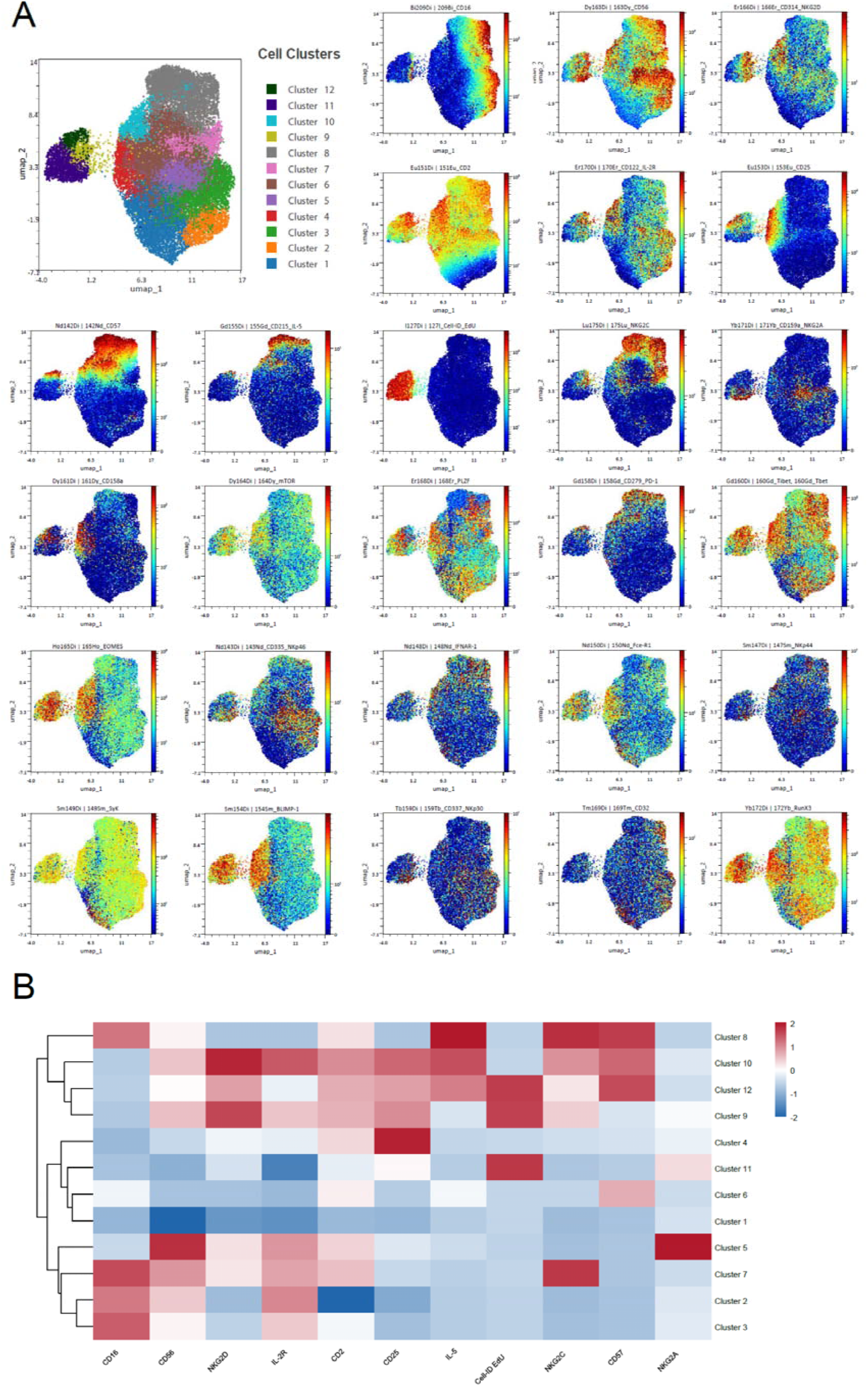
Mass Cytometry Cluster analyses. **(A)** Expression of all markers included in the CyTOF panel is displayed across the identified cell clusters using UMAP. The following markers were used for FlowSOM clustering: CD16, CD56, NKG2D, IL-2R/CD122, CD2, CD25, IL-5, Cell-ID EdU, NKG2C, CD57, and NKG2A. **(B)** Heatmap showing the z-scored expression of the markers used for clustering across the identified cell clusters. The row dendrogram illustrates similarities between clusters based on their overall marker expression profiles, with more closely related clusters grouped together.

**Fig. S8.**
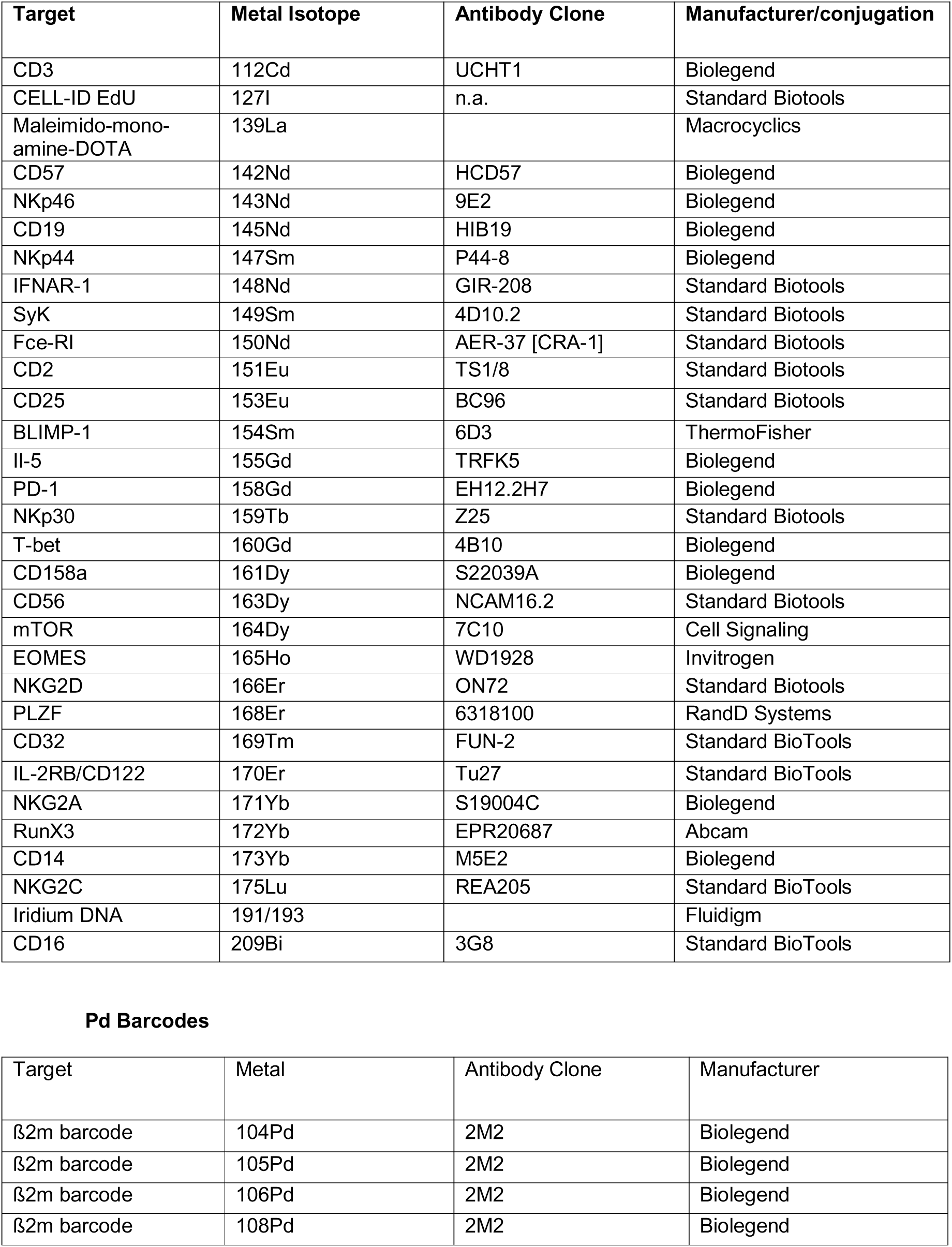
**Mass Cytometry panel**

